# The insulating activity of the *Drosophila* BX-C chromatin boundary *Fub-1* is parasegmentally regulated by lncRNA read-through

**DOI:** 10.1101/2022.11.13.516321

**Authors:** Airat Ibragimov, Xin Yang Bing, Yulii Shidlovskii, Mike Levine, Pavel Georgiev, Paul Schedl

## Abstract

Though long non-coding RNAs (lncRNAs) represent a substantial fraction of the Pol II transcripts in multicellular animals, only a few have known functions. Here we report that the blocking activity of the Bithorax complex (BX-C) *Fub-1* boundary is segmentally regulated by its own lncRNA. The *Fub-1* boundary is located between the *Ultrabithorax* (*Ubx*) gene and the *bxd/pbx* regulatory domain, which is responsible for regulating *Ubx* expression in parasegment PS6/segment A1. *Fub-1* consists of two hypersensitive sites, *HS1* and *HS2. HS1* is an insulator while *HS2* functions primarily as a lncRNA promoter. To activate *Ubx* expression in PS6/A1 enhancers in the *bxd/pbx* domain must be able to bypass *Fub-1* blocking activity. We show that expression of the *Fub-1* lncRNAs in PS6/A1 from the *HS2* promoter inactivates *Fub-1* insulating activity. Inactivation is due to readthrough as the *HS2* promoter must be directed towards *HS1* to disrupt blocking.

## INTRODUCTION

The vast majority of Pol II transcripts encoded by the genomes of multicellular animals correspond to long non-coding RNAs (lncRNAs), not mRNAs. Their wide distribution in the genome and the fact that their expression is often coordinated with genes in the immediate vicinity has led to the idea that they have functions in gene regulation and chromosome organization (Herman et al., 2022; Li and Fu, 2019; Núñez-Martínez and Recillas-Targa, 2022; Statello et al., 2021). Unlike the sequences of protein coding mRNAs, which are typically conserved across species, the primary sequences of lncRNAs are usually not very well conserved. However, some lncRNAs contain short sequence blocks that exhibit a relatively high degree of conservation, even though most of the lncRNA is divergent. In these instances, the lncRNA itself has regulatory activities. A classic example is the mammalian X chromosome inactivation lncRNA Xist. Conserved RNA sequence blocks within the Xist transcript function in recruiting Polycomb (PcG) silencing factors and targeting the transcript to the X chromosome (Jacobson et al., 2022; Lu et al., 2020; Pandya-Jones et al., 2020). Other lncRNAs, like the HoxA complex lncRNA *HOTT1P* encode RNA sequences that recruit chromatin modifiers that help promote gene expression (Wang et al., 2011). In other cases, the lncRNA itself does not appear to have important functions. Instead, it is the regulatory elements (enhancers, silencers and promoters) that control the expression of the lncRNA that are functionally important. For example, the expression of the Myc gene is down regulated by a mechanism in which the promoter for the *PVT1* lncRNA competes with Myc for physical interactions with a set of shared enhancers. In other cases, the transcription of the lncRNA appears to be the important regulatory function. Isoda et al. found that transcription of *ThymoD* lncRNA helps regulate the expression of Bcl11b, a gene that plays an important role in specifying T cell fate (Isoda et al., 2017). Transcription of the lncRNA induces the demethylation of recognition sites for the chromosomal architectural protein CTCF, enabling CTCF to bind to these sites and induce looping between the Bcl11b enhancers and the Bcl11b gene.

In the studies reported here we have investigated the role of lncRNAs in regulating the chromatin organization and expression of the homeotic gene, *Ultrabithorax* (*Ubx*), in the *Drosophila* bithorax complex (BX-C). *Ubx* together with *abdominal-A* (*abd-a*) and *Abdominal-B* (*Abd-B*) are responsible for specifying the identity of the nine parasegments (PS)/segments that form the posterior 2/3rds of the fly ((Maeda and Karch, 2015): Fig. 1). Parasegment identity depends upon a series of nine regulatory domains that direct the temporal and tissue specific patterns of expression of the three BX-C homeotic genes. *Ubx* is responsible for specifying parasegments PS5 (adult cuticle segment T3) and PS6 (adult cuticle segment A1). Its expression in these two parasegments is controlled by the *abx/bx* and *bxd/pbx* regulatory domains respectively. *abd-a* expression in PS7-PS9 is directed by the *iab-2--iab-4* regulatory domains, while *Abd-B* expression in PS10-14 is regulated by the *iab-5--iab-9* regulatory domains. Each regulatory domain has an initiation element, a set of tissue-specific enhancers, and Polycomb Response Elements (PREs) (Iampietro et al., 2010; Maeda and Karch, 2015, 2006). Early in development, a combination of maternal, gap and pair-rule gene proteins interact with the initiation elements in each regulatory domain, and set the domain in either the *“off”* or *“on ”* state. The BX-C regulatory domains are sequentially activated along the anterior-posterior axis of the embryo. For example, in PS5(T3) *abx/bx* is activated, while the remaining BX-C domains including *bxd/pbx* are *“off”*, and it is responsible for regulating *Ubx* expression in this parasegment. Both *abx/bx* and *bxd/pbx* are turned on in PS6(A1); however, *Ubx* expression, and parasegment identity depend on the *bxd/pbx* domain. Once the activity state of the domain is set in early embryos, it is remembered during the remainder of development by the action of Polycomb Group proteins (PcG: *off*) and trithorax group proteins (Trx: *on*). Each domain also contains tissue and stage specific enhancers which drive the expression of its target homeotic gene in a pattern appropriate for proper development of the parasgement it specifies.

**Figure.1.**
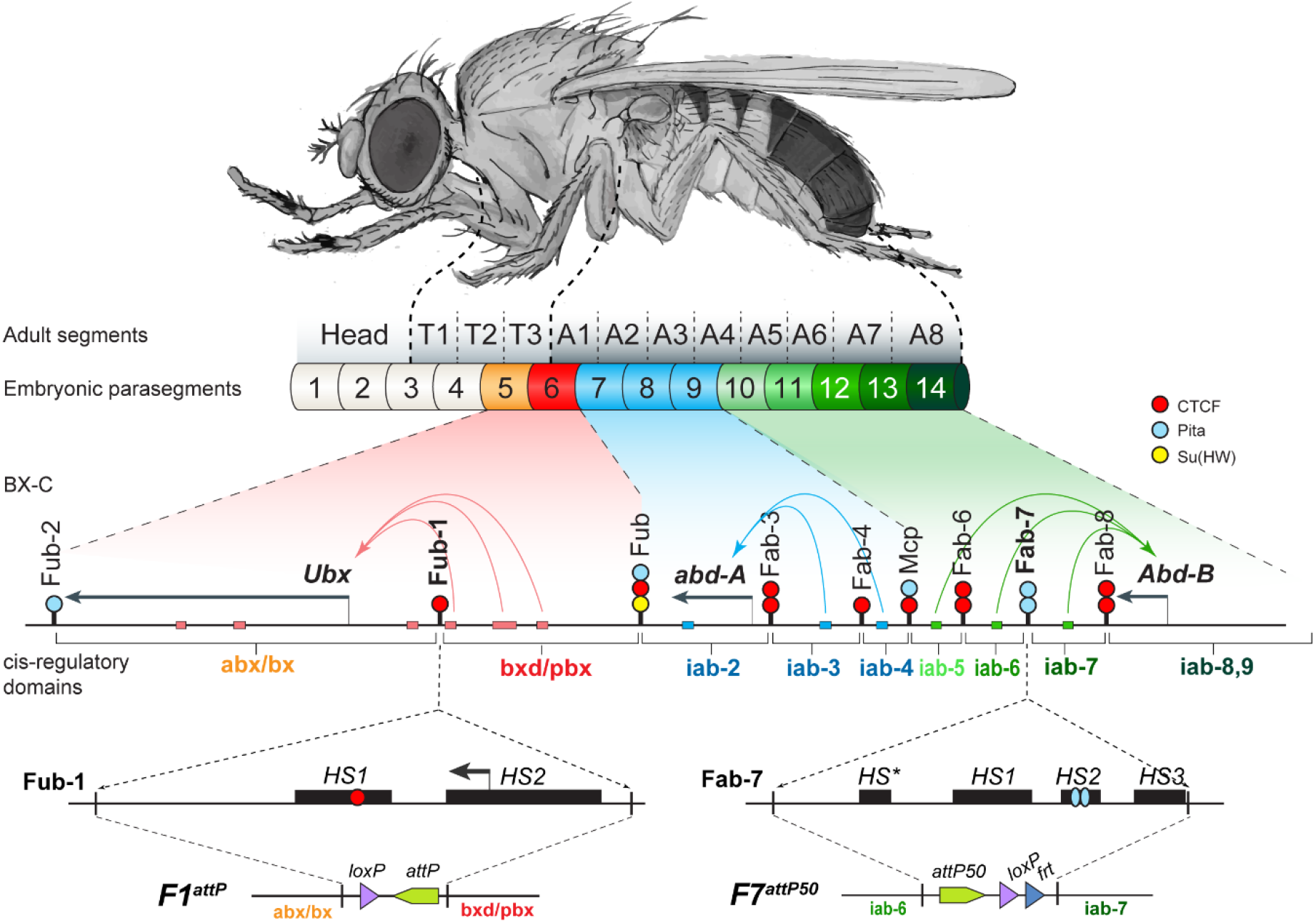
The organization of the genes and regulatory domains in BX-C. The *Drosophila melanogaster* Bithorax complex (BX-C), which contains the Hox genes *Ubx*, *abd-A* and *Abd-B*, is shown in relation to where these three genes are expressed in an embryo. There are nine *cis*-regulatory domains that are responsible for regulation of *Ubx* (*abx/bx* and *bxd/pbx* domains), *abd-A* (*iab-2-4* domains) and *Abd-B* (*iab-5–7* and *iab-8,9* domains), and for the development of parasegments 5 to 13 (PS)/segments (T3-A8). The anterior limit of expression of the three Hox gene is indicated by color coding: red: *Ubx*; blue: *abd-A*; green: *Abd-B* (reviewed in (Maeda and Karch, 2015)). The lines with colored circles mark chromatin boundaries. dCTCF, Pita, and Su(Hw) binding sites at the boundaries are shown as red, blue, and yellow circles/ ovals, respectively. Embryonic enhancers are indicated by pink, blue and green bars on coordinate line. On the bottom of the figure, the molecular maps of the *Fub-1* and *Fab-7* boundaries are shown, including their deletions. Transposase/nuclease hypersensitive sites are shown as black boxes above the coordinate bar. The proximal and distal deficiency endpoints of the *Fub-1* and *Fab-7* deletions used in the replacement experiments are indicated by vertical lines. The *attP*, *lox*, and *frt* sites used in genome manipulations are shown as green, violet, and blue triangles, respectively.

In order to properly specify parasegment identity, the nine BX-C regulatory domains must be functionally autonomous. Autonomy is conferred by boundary elements (insulators) that flank each domain (Maeda and Karch, 2015). The role of boundaries is BX-C regulation is best understood in the *Abd-B* region of the complex. Mutations that inactivate *Abd-B* boundaries cause a gain-of-function (GOF) transformation in segment identity. For example, the *Fab-7* boundary separates the *iab-6* and *iab-7* regulatory domains which are responsible for specifying PS11(A6) and PS12 (A7) identity respectively (Gyurkovics et al., 1990). When *Fab-7* is deleted, the *iab-6* initiator activates the *iab-7* domain in PS11(A6) and *iab-7* instead of *iab-6* drives *Abd-B* expression in this parasegment as well as in PS12(A7). This results in the duplication of the PS12/A7 segment. However, blocking crosstalk between adjacent regulatory domains is not the only function of BX-C boundaries. The three homeotic genes and their associated regulatory domains are arranged so that all but three of the domains are separated from their targets by one or more intervening boundaries. For this reason, all but two (*Fub* and *Mcp*) of the internal BX-C boundaries must also have bypass activity (Hogga et al., 2001; Kyrchanova et al., 2019b, 2019a, 2015). For example, *Fab-7* has bypass activity and it mediate regulatory interactions between *iab-6* and *Abd-B*. However, when *Fab-7* is replaced by generic fly boundaries (e.g., *scs* or *su(Hw)*) or by BX-C boundaries that lack bypass activity (*Mcp*) these boundaries block crosstalk but do not support bypass (Hogga et al., 2001; Kyrchanova et al., 2017). As a consequence, *Abd-B* expression in PS11/A6 is driven by *iab-5* not *iab-6* and this results in a loss-of-function phenotype.

In the *Ubx* region of the complex, the *bxd/pbx* regulatory domain is separated from the *Ubx* gene by a boundary, *Fub-1* (Fig. 1) (Bowman et al., 2014, p. 27; Mateo et al., 2019). This boundary would have to be bypassed so that *bxd/pbx* can direct *Ubx* expression in PS5/A1. Unlike the *Abd-B* region of the complex where multiple boundaries are located between the *iab-5* and *iab-6* regulatory domain, there is only one boundary between *bxd/pbx* and the *Ubx* gene. As a consequence, the bypass mechanism might differ from that deployed elsewhere in BX-C. Here we show that segmentally regulated transcription of a lncRNA disrupts the blocking activity of *Fub-1*, enabling enhancers in the *bxd/pbx* regulatory domain to activated *Ubx* in PS6/A1 cells.

## RESULTS

### The *bxd/pbx* regulatory domain: boundaries and lncRNAs

The *bxd/pbx* regulatory domain controls *Ubx* expression in A1/PS6. This domain is *off* in more anterior parasegments (segments), while it is turned on in A1/PS6 and more posterior parasegments. The *bxd/pbx* domain is bracketed by the *Fub* boundary on the centromere distal side and *Fub-1* on the centromere proximal side. The chromosomal architectural proteins CTCF, Pita and Su(Hw) are associated with *Fub* in ChIP experiments, and like boundaries the *Abd-B* region of BX-C *Fub* deletions result in a GOF transformation of PS6(A1) into PS7(A2). The *Fub-1* boundary has only been defined molecularly. ATAC-seq mapping in nuclear cycle 14 embryos indicate that this boundary includes two chromatin-specific transposase hypersensitive regions: *HS1* ~ 200 bp and *HS2* ~ 300 bp (Figure 1) (Hannon et al., 2017). In ChIP experiments *Fub-1* is bound by the chromosomal architectural proteins CTCF, CP190, Zw5, Pita, and Elba (Bowman et al., 2014, p. 27; Ueberschär et al., 2019; Zolotarev et al., 2016) (Sup Fig.2). In addition, RNA Pol II and the histone mark for active transcription, H3K4me3 both map to *Fub-1.* Consistent with results described below, the signal for H3K4me3 in PS7 cell ChIPs is greater than that observed in mixed ChIPs (Bowman et al., 2014, p. 27). In between *Fub-1* and the *Ubx* promoter there are two prominent Zelda peaks and a putative enhancer (Sup Fig.2) (Harrison et al., 2011).

**Figure.2.**
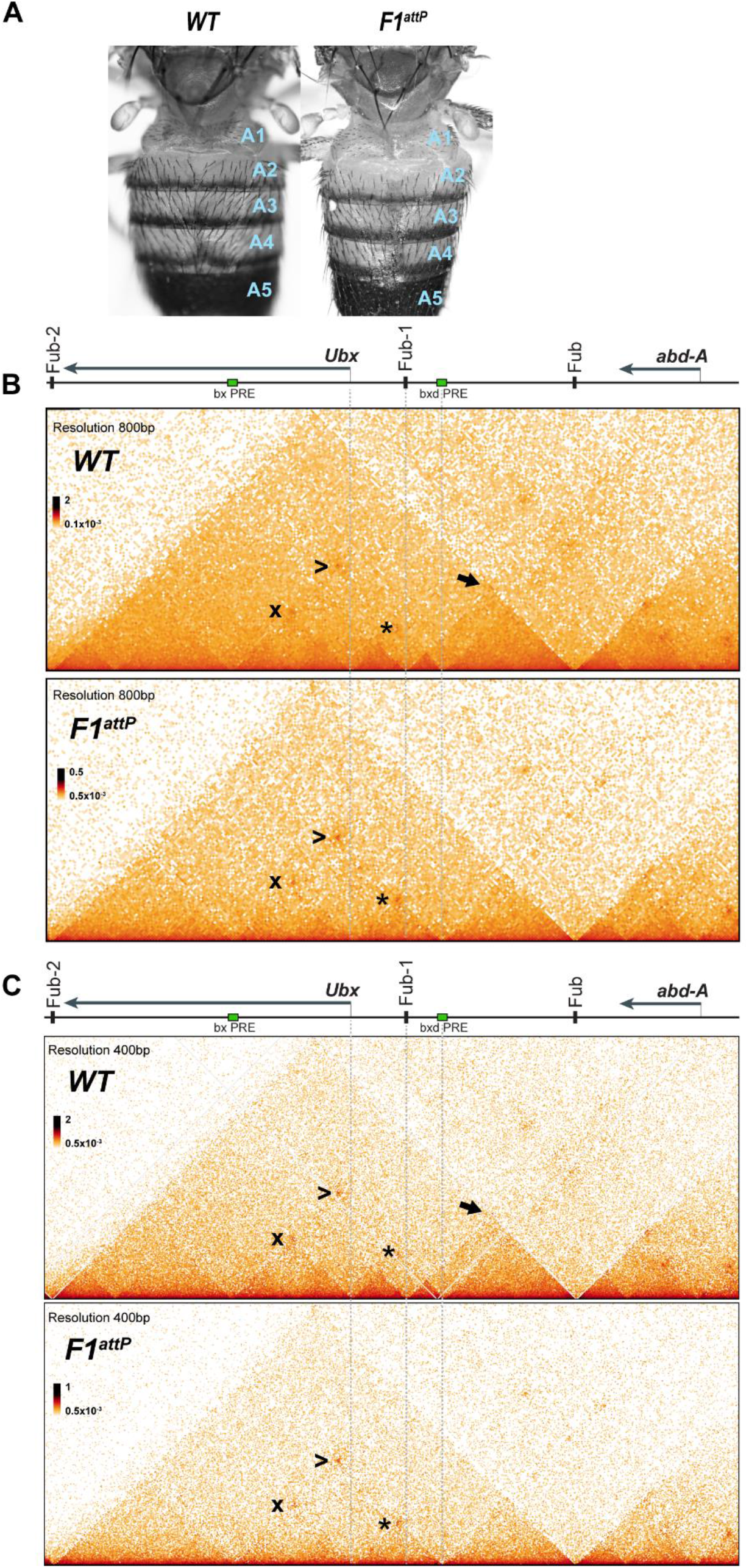
Chromatin topology in *bxd/pbx* domain in *WT* and in *Fub-1* deletion. (A) Morphology of the abdominal segments of *WT* and *F1^attP^* flies. (B) Micro-C contact map of the *WT* and *F1^attP^* embryos at 800 bp resolution. The black arrow points to a sub-TAD formed by *Fub-1* and *Fub* boundaries. The *Ubx* promoter is linked to the *bxd*. PRE by a LDIC, that is marked at the apex by an interaction dot (*). The *bxd* PRE forms an interaction dot (>) with the *bx* PRE as does the *Ubx* promoter (x). (C) Micro-C contact map of the *WT* and *F1^attP^* embryos at 400 bp resolution.

Besides directing the expression of *Ubx* in PS6(A1) and more posterior parasegements, the *bxd/pbx* domain also controls the expression of two lncRNAs, *bxd* and *F1^HS2^* (Fig 5D). Peace et al found that the *bxd* lncRNA initiates from a promoter close to the *pbx* enhancer and extends into the region located in between *Fub-1* and the *Ubx* promoter (Pease et al., 2013). The *bxd* lncRNA is first detected at the blastoderm stage as a broad band that extends from PS5/A1 to PS12 (A7). Between the onset of gastrulation and stage 10 *bxd* lncRNA expression resolves into a series of 8 stripes of differing intensity that span parasegments PS5-PS12. The *bxd* lncRNA then disappears when germband retraction commences. Peace et al also detected a second *bxd/pbx* dependent lncRNA, *F1^HS2^*. Like *bxd* lncRNA it is expressed in PS6 (A1) and more posterior parasegments. This lncRNA is encoded by sequences located between *Ubx* and *Fub-1* including sequences in *Fub-1 HS1* and based on the *in situ* experiments of Peace et al., it likely originates from *HS2*. As *F1^HS2^* shares exon sequences with the *bxd* lncRNA, it is not clear whether it is expressed in the period between the blastoderm stage and the onset of germband retraction. However, the *F1^HS2^* lncRNA is detected in PS6/A1 and more posterior segments after the *bxd* lncRNA disappears at the onset of germband retraction.

**Figure.3.**
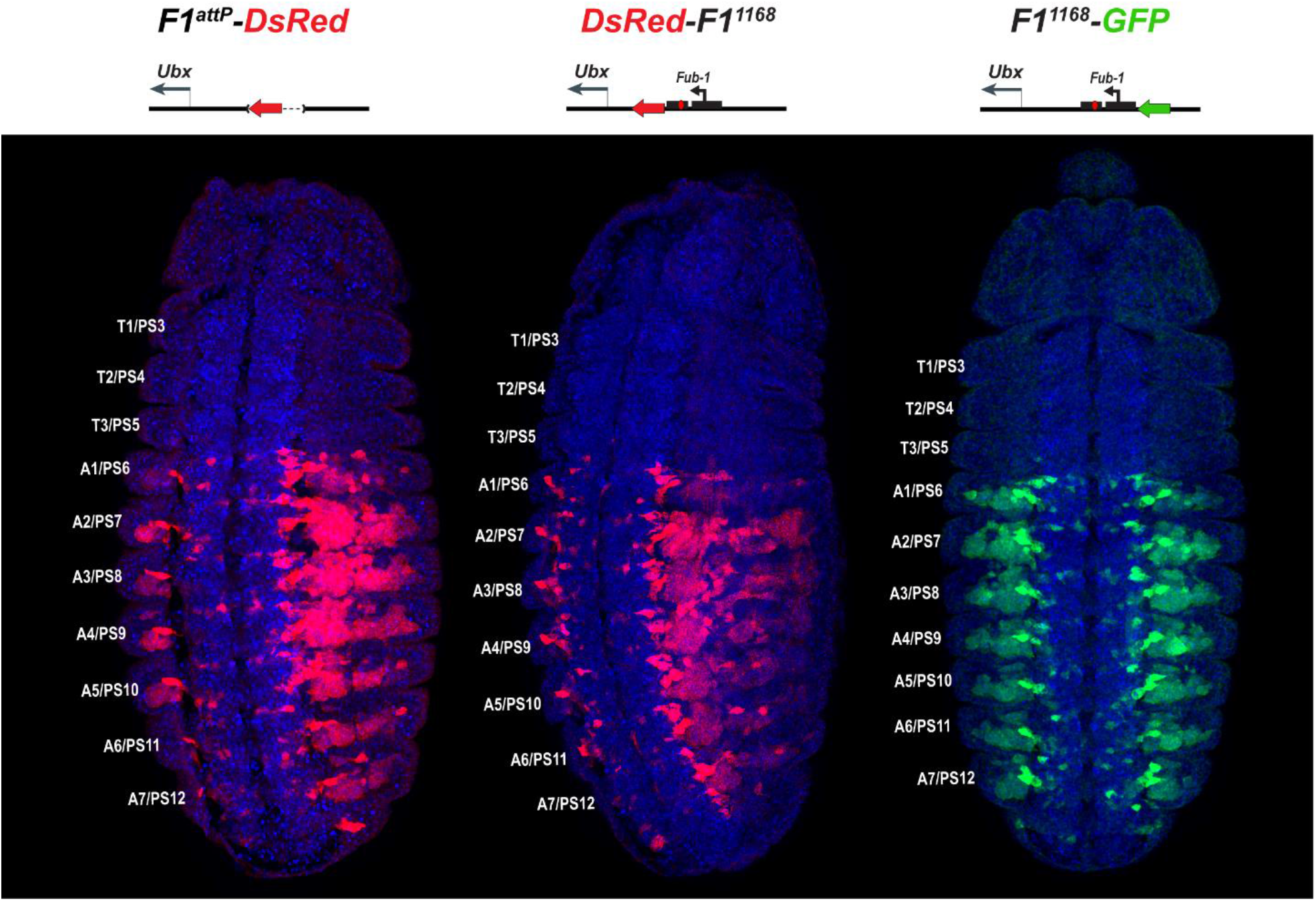
*DsRed* and *GFP* markers trap enhancer activity in *Fub-1* replacements. *DsRed* (in red) and *GFP* (in green) expression in stage 14 embryos in the *Fub-1* deletion and in two replacements as indicated. Hoechst was used to stain nuclei (in blue) *DsRed* expression in *F1^attP^-DsRed* begins in PS6/A1. In *DsRed-F1^1168^* and *F1^1168^-GFP* expression patterns of both markers are limited to PS6/A1-PS13/A8, consistent with the idea that *Ubx* promoter region demarcate *abx/bx* and *bxd/pbx* domains.

**Figure 4.**
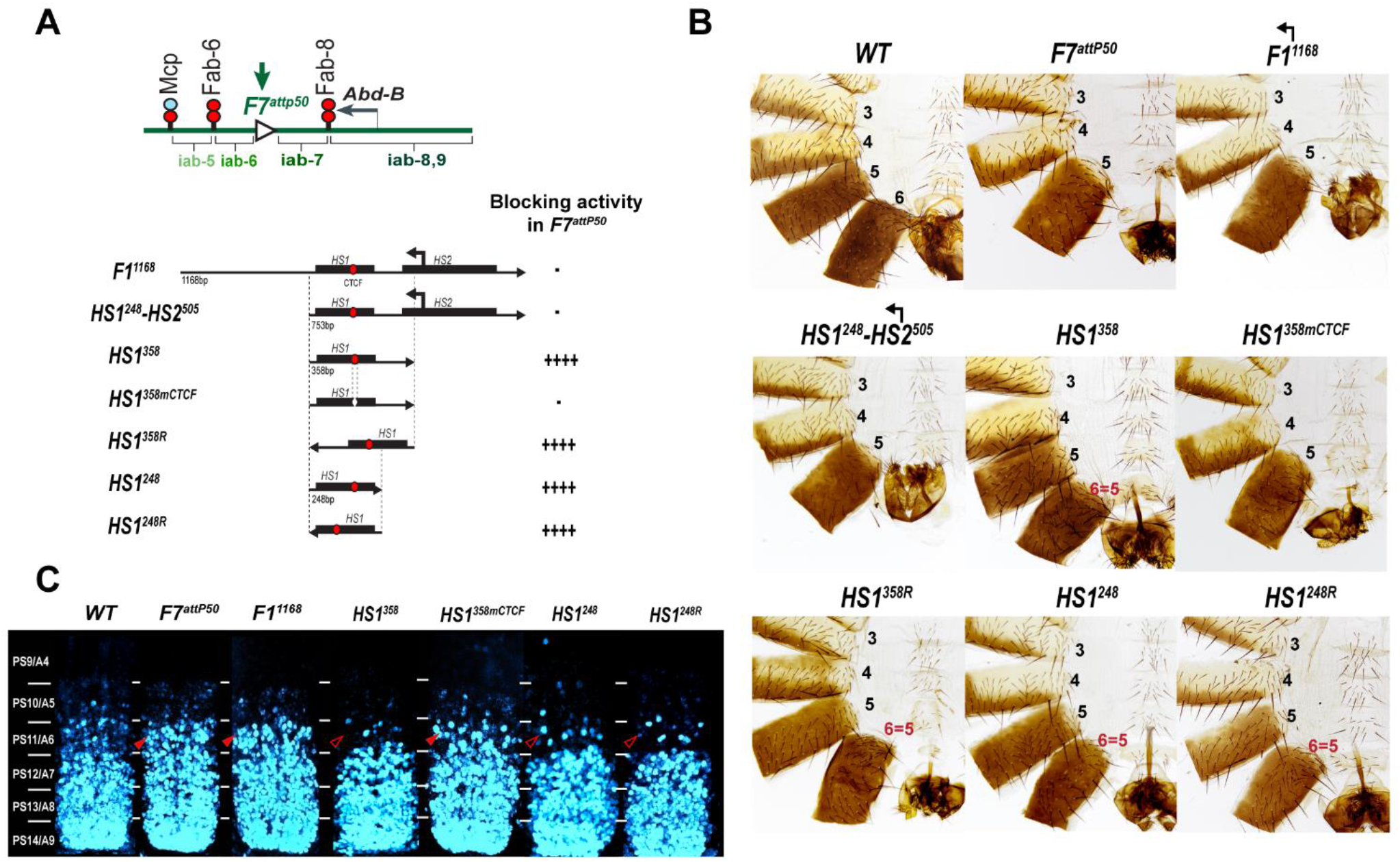
Testing boundary activity of *Fub-1* sequences. (A) Top: Schematic presentation of *Fab-7* substitution. Bottom: *Fub-1* fragments used in the replacement experiments. On the right side, the insulator activity with the various fragments in the *F7^attP50^* insertion site in adults as judged from cuticle preps. The number of “+” signs reflects the strength of insulator activity, where “++++” is full blocking, and “-” is lack of detectable blocking activity, respectively. Designations are the same as described in Figure 1. (B) Morphology of the male abdominal segments (numbered) in *F1^1168^, HS1^248^-HS2^505^, HS1^358^, HS1^358mCTCF^, HS1^358R^, HS1^248^* and *HS1^248R^* replacements. (C) *Abd-B* expression in Fab-7 replacement embryos. Each panel shows a confocal image of the embryonic CNS of stage 15 embryos stained with antibodies against *Abd-B*. (cyan). The filled red arrowheads show morphological features indicative of GOF transformations. The empty red arrowheads show the signs of the LOF transformation, which is directly correlated with the boundary function of tested DNA fragments. The *WT* expression pattern of *Abd-B* in the embryonic CNS is characterized by a stepwise gradient of increasing protein level from PS10/A5 to PS14/A8. In *F7^attP50^* embryos *Abd-B* expression level in PS11/A6 is roughly equal to that in PS12/A7, indicating *iab-7* drives *Abd-B* expression in PS11/A6 (GOF phenotype). Consistent with the adult phenotype, in *F1^1168^* and *HS1^248^-HS2^505^ Abd-B* expression in PS11/A6 is the same as in *F7^attP50^* The *Abd-B* expression pattern in *HS1^358^* and *HS1^248^* replacement is also consistent with the adult cuticular phenotypes: *Abd-B* expression is reduced in both PS10/A5 and PS11/A6 (LOF phenotype) compared with *WT*. In contrast, mutation of dCTCF site in *HS1^358mCTCF^* results in the loss of blocking activity and *Abd-B* expression pattern similar to *F7^attP50^*.

**Figure 5.**
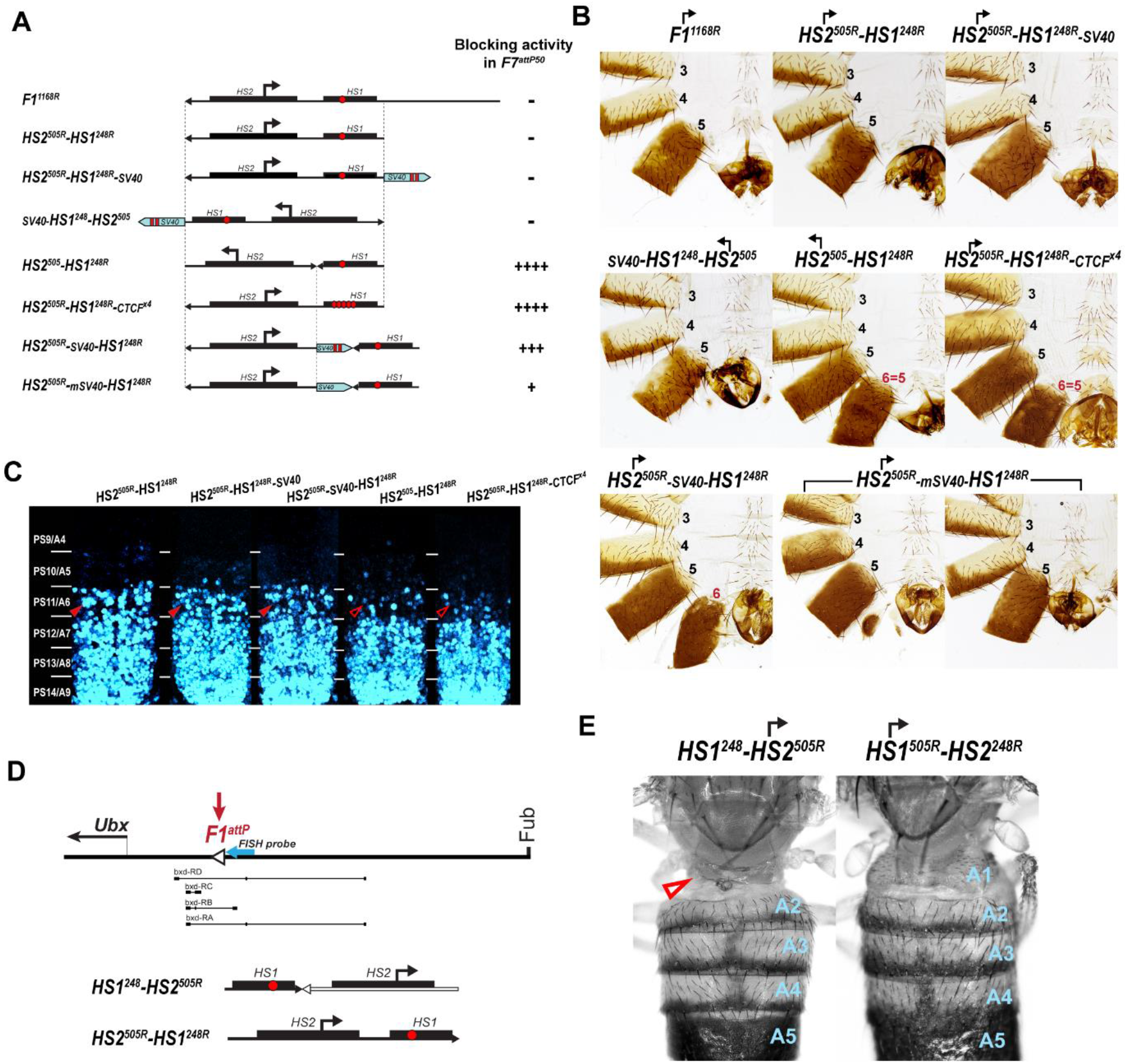
Read-through transcription inactivates *HS1* insulator. (A) Schematic representation of sequences tested in *F7^attP50^.* SV40 terminator is shown as a light blue arrow, poly(A) sites are marked as vertical red lines. All other designations are the same as described in Figure 4. (B) Morphology of the male abdominal segments (numbered) of *Fab-7* replacements. (C) *Abd-B* expression in CNS of *Fab-7* replacement embryos. Embryos were stained and marked as in Fig 4. (D) Schematic presentation of *F1^attP^* landing platform with *HS1^248^-HS2^505R^* and *HS2^505R^-HS1^248R^* replacements. All designations are the same as described in Figure 1. Characterized *bxd* lncRNA transcripts are presented under coordinate line. The smFISH strand-specific probes are shown as one light blue arrow (see Table S2 for sequences). (E) Morphology of the abdominal segments of *HS1^248^-HS2^505R^* and *HS2^505R^-HS1^248R^* flies. The red arrow shows the signs of the LOF phenotype: complete reduction of A1 segment and the appearance of postnotal tissue on its place.

As has been reported for the regulatory elements associated with other lncRNAs, the *Fub-1* hypersensitive sites are evolutionarily conserved. As shown in Sup Fig.3, key sequence blocks within *HS1* and *HS2* are detected not only in other members of the *Sophophora* subgenus, but also in *D. virilis* a distantly related member of the *Drosophila* subgenus. The conserved blocks in *HS1* include the dCTCF recognition site, while in *HS2* the conserved blocks include several GAGA motifs which correspond to binding sites for the GAGA factor (GAF) as well as the predicted transcription start sites for the *F1^HS2^* lncRNA.

### *Fub-1* subdivides the *Ubx* regulatory domains

These observations, together with the studies of Bowman et al. which showed that *Fub-1* marks the border of a Polycomb histone H3K27me3 domain in PS5 cells would predict that *Fub-1* delimits the endpoint of two adjacent TADs. The centromere distal TAD would encompass the *bxd/pbx* regulatory domain and end at *Fub* boundary which is located downstream of the *abd-A* transcription unit (Bender and Lucas, 2013; Bowman et al., 2014, p. 27). The centromere proximal TAD would encompass the entire *Ubx* transcription unit and end at a predicted Pita/CP190 boundary, *Fub-2*, that is located between the 3’ end of the *Ubx* gene and the 5’ end of *modSP*.

To test these predictions, we used Micro-C to examine the TAD organization of *Ubx* and its two regulatory domains in 12-18h embryos. As shown in Fig. 2 both regulatory domains (plus the *Ubx* transcription unit) are encompassed in a large TAD that extends from *Fub-2* on the centromere proximal side of *Ubx* to *Fub* on the distal side. Within this large TAD, there is a ~50 kb sub-TAD which spans *bxd/pbx* and is delimited by *Fub-1* on the left and *Fub* on the right (Fig.2 arrow). This *bxd/pbx* sub-TAD includes several smaller chromosomal segments that exhibit a high density of internal contacts (HDIC). The endpoint for one of these HDIC domains maps to *Fub-1*, while the other endpoint maps to the *bxd* PRE. There is a second HDIC domain linking *Fub-1* to sequences just upstream of the *Ubx* promoter. The *Ubx* promoter is also linked to the *bxd* PRE by a lower density of internal contacts domain (LDIC), that is marked at the apex by an interaction dot (*). The *bxd* PRE also forms an interaction dot (>) with the *bx* PRE as does the *Ubx* promoter (x). Interestingly PREs in other developmental loci have recently been shown to interact with each other and also function as “tethering” elements helping to link distant enhancers to their target genes (Batut et al., 2022; Eagen et al., 2017; Levo et al., 2022).

### *Fub-1* is not required to block crosstalk between *abx/bx* and *bxd/pbx*

A BX-C boundary in the location of *Fub-1* would be expected to have two regulatory functions. One would be blocking crosstalk between *abx/bx* and *bxd/pbx* while the other would be bypass so that enhancers in the *bxd/pbx* domain would be able to drive *Ubx* expression in PS6/A1. To test for blocking activity, we generated a 1168-bp deletion (*F1^attP^*) that retains an *attP* site for boundary replacement experiments (Figure S1A). Deletions of boundaries elsewhere in BX-C typically result in a GOF transformations. This transformation arises because initiators in the domain centromere proximal (right: Fig. 1) to the boundary are able to active the domain distal to the boundary (left: Fig. 1) in a parasegment in which the distal domain would normally be silent (Barges et al., 2000; Bender and Lucas, 2013; Gyurkovics et al., 1990; Karch et al., 1994; Postika et al., 2021). For this reason, we anticipated that the *bxd/pbx* domain would be inappropriately activated by initiators in *abx/bx* in PS5, and in adults we would observe a GOF transformation of T3 (PS5) towards A1 (PS6). Unexpectedly, however, *F1^attP^* flies do not show any evidence of a GOF transformation and their morphology is indistinguishable from wild type (*WT*) (Fig. 2A).

A plausible explanation for this result is that there is a nearby element that can substitute for *Fub-1* and block crosstalk between *abx/bx* and *bxd/pbx*. Fig. 2B,C show that the sub-TAD linking *Fub-1* to *Fub* that encompassed the *bxd/pbx* regulatory domain disappears in the *Fub-1* deletion. However, a new sub-TAD that includes *Fub* does not appear to be formed. Instead, the element located just upstream of the *Ubx* promoter forms a sub-TAD domain with the *bxd* PRE (see interaction dot (Fig 2B,C *) at apex). It would appear that this topological configuration is sufficient to block crosstalk between initiators in *abx/bx* and *bxd/pbx* when the activity state of these domains in T3 (PS5) is set during early embryogenesis.

### The *Fub-1* element does not block enhancers in *bxd/pbx* from activating reporters

Although *Fub-1* defines one end-point of a TAD and marks the border for the PcG H3K27me3 histone mark in PS5 nuclei, the finding that *Fub-1* deletions have no phenotype raises the possibility that it is actually “within” the *bxd/pbx* regulatory domain. If this is the case then reporters inserted between *Fub-1* and the element just upstream of the *Ubx* promoter should respond to *bxd/pbx* enhancers, but be insulated from enhancers in the *abx/bx* regulatory domain. To test this possibility, we analyzed the pattern of expression of reporters inserted into the *F1^attP^* site with and without the *Fub-1* boundary.

We first examined the *DsRed* reporter used to mark the *Fub-1* deletion in *DsRed-F1attP*. Fig. 3 shows that *DsRed* is expressed throughout the posterior ~2/3rds of *Fub-1* deletion embryos, with an anterior limit corresponding to the anterior border of PS6/A1. Thus, as expected from the lack of phenotypic effects of the *Fub-1* deletion, the reporter is insulated from the *abx/bx* domain, but is subject to regulation by the *bxd/pbx* domain. We next introduced the 1168-bp *Fub-1* fragment together with a *GFP* reporter flanked by *frt* sites back into the *DsRed-F1^attP^* platform (Figure S1). In the resulting insert the *Fub-1* element is flanked by *DsRed* on the proximal *Ubx* side, and *GFP* on the distal *bxd/pbx* side. The reporters were then excised individually to give *DsRed-Fub-1* and *Fub-1-GFP* (Figure S1). As shown in Fig. 3, the anterior border of expression of both *DsRed* and *GFP* corresponds to the PS6/A1 as is observed for *DsRed* when the *Fub-1* element is deleted. Thus *Fub-1* is unable to block the *bxd/pbx* enhancers from activating the *DsRed* reporter in PS6/A1 cells (and more posterior parasegments/segments) and this reporter is also “located” within the *bxd/pbx* regulatory domain.

These findings fit with previous enhancer trap studies which showed that P-element insertions around the *Ubx* promoter displayed two distinct patterns of expression. An enhancer trap located 13-bp upstream of the *Ubx* transcription start site had a PS5/T3 anterior border indicating that it is regulated by the *abx/bx* domain. In contrast, an enhancer trap located 196 bp upstream of the start site had an anterior limit of expression shifted to PS6/A1 (McCall et al., 1994). Taken together with the experiments described in the previous section, it would appear that the *Ubx* promoter region is sufficient for the functional autonomy of the *abx/bx* and *bxd/pbx* regulatory domains under laboratory growth conditions.

### *Fub-1* replacements of *Fab-7* do not block crosstalk between *iab-6* and *iab-7*

The results described above are unexpected for several reasons. First, *Fub-1* defines the proximal endpoint of a TAD that extends to *Fub* and encompasses the *bxd/pbx* regulatory domain. Second it also corresponds to the proximal border of the PcG H3K27me3 histone mark in PS5 nuclei where *abx/bx* should be *“on ”* and *bxd/pbx* should be *“off”* Third, both of the *Fub-1* sub-elements, *HS1* and *HS2*, are evolutionarily conserved. In spite of these properties, deletion of *Fub-1* does not result in a detectable GOF transformation of T3/PS5 into A1/PS6. This is likely explained by the presence of the boundary element upstream of the *Ubx* promoter which is able to block crosstalk between initiators in *abx/bx* and *pbx/bxd*. However, this would not explain why *Fub-1* fails to block the *pbx/bxd* enhancers from activating a *DsRed* reporter when interposed between the enhancers and the reporter. These observations raise a novel possibility: even though *Fub-1* has many of the characteristic chromosome architectural functions of boundary elements it differs from other boundaries in flies and other organisms in that it lacks insulating activity.

To test whether *Fub-1* lacks insulating activity we took advantage of the *Fab-7^attP50^* (*F7^attP50^*) replacement platform (Wolle et al., 2015)(Figure 1). In this platform, the *Fab-7* boundary was deleted, and an *attP* site introduced in its place. The deletion of *Fab-7* fuses the *iab-6* and *iab-7* regulatory domains, enabling parasegment-specific initiation elements in *iab-6* to ectopically activate *iab-7* (Karch et al., 1994; Wolle et al., 2015). As a consequence, *iab-7* drives *Abd-B* expression in PS11, transforming PS11/A6 into a duplicate copy of PS12/A7 (Figure 1). The *attP* site in the *F7^attP50^* can be used to insert any sequences of interest to test for two different boundary functions. The first is blocking crosstalk between *iab-6* and *iab-7*, while the second is supporting bypass so that *iab-6* can regulated *Abd-B* in PS11/A6.

In the first experiment, we used the *F7^attP50^* replacement platform to test the blocking activity of a 1168-bp sequence spanning entire deletion, *Fub-1^1168^*, and a 753-bp sequence including only hypersensitive sites, *HS1^248^-HS2^505^* (R6.22: 3R:16748578-16749330) inserted in the same 5’→3’ orientation as the endogenous *Fub-1* element. Figure 4B and S4 show that males carrying either the 1168 bp or the 753 bp replacements lack A6 just like the starting *F7^attP50^* deletion. This result indicates that these two *Fub-1* fragments do not prevent cross-talk between *iab-6* and *iab-7*. The *HS1^248^-HS2^505^* replacement gain-of-function (GOF) phenotype is reflected in the pattern of *Abd-B* expression in the embryonic CNS which closely matches that of the starting *F7^attP50^* deletion (Fig 4C). We also inserted the two *Fub-1* fragments in the reverse orientation; however, in this case also neither of the fragments rescued the GOF phenotype of the *F7^attP50^* deletion (Fig. 5).

### *HS1* has boundary activity

Previous replacement experiments have shown that dCTCF associated BX-C boundaries like *Mcp* are able to block crosstalk between *iab-6* and *iab-7* (Postika et al., 2018). For this reason, we wondered whether *HS1* alone has blocking activity. We used two different *HS1* fragments to test this possibility. One was 358 bp (R6.22: 3R: 16748578-16748935) and included all of the *HS1* sequences plus the sequences located between *HS1* and *HS2*. The other, a 248 bp fragment (R6.22: 3R: 16748578-16748825), contains only the *HS1* region. These fragments were inserted in both the forward and reverse orientations. We also tested a 505 bp fragment spanning the *HS2* (R6.22: 3R: 16748826-16749330) in both the forward and reverse orientations (Sup Fig. 4,5). While *HS2* lacks blocking activity (Sup Fig. 4,5), Fig. 4B shows that both of the *HS1* fragments block crosstalk between *iab-6* and *iab-7* and rescue the GOF transformation of A6 into A7. The rescuing activity is orientation independent. In all four cases, however, the morphology of the A6 segment resembles that normally observed in A5, not A6. This LOF phenotype is observed when the *Fab-7* replacements have insulating activity but cannot support bypass (Hogga et al., 2001; Kyrchanova et al., 2017). Consistent with this conclusion, *Abd-B* expression in the embryonic CNS is suppressed in A5 and A6 by the *HS1* fragments (Fig. 4C).

dCTCF recognition sequences were found to be required for the blocking activity of other CTCF associated BX-C boundaries (Kyrchanova et al., 2017, 2016, p. 8). This is also true for the dCTCF site in *HS1*. Fig. 4 shows that a mutation in the *HS1* dCTCF binding site completely disrupts insulating activity. Like the starting *F7^attP50^* platform, the adult *HS1^358mCTCF^* males lack the A6 segment indicating that PS11/A6 is fully transformed into a copy of PS12/A7 (Figure 4B and S4). Consistent with a loss of blocking activity, the pattern of *Abd-B* expression in *HS1^358mCTCF^* embryos is similar to that of *F7^attP50^* (Figure 4C).

### Read-through transcription from *HS2* abrogates *HS1* insulator function

The finding that fragments containing only *HS1* have insulating activity, while the full-length *Fub-1* fragment does not, suggests that when *HS2* is present it inactivates the *HS1* boundary. One likely mechanism is transcriptional read-through. Studies on the induction of the chicken lysozyme gene by bacterial lipopolysaccharides (LPS) by Lefevre et al. [36], showed that LPS treatment activated the transcription of a lncRNA through a CTCF boundary (Lefevre et al., 2008). Read-through transcription resulted in the inactivation of the boundary enabling distal enhancers to bypass the boundary and turn on the lysozyme gene.

To test this hypothesis, we inserted a 229-bp SV40 transcription terminator in between *HS2* and *HS1* to give the *HS2^505R^-SV40-HS1^248R^* replacement (Figure 5A). Unlike the starting *HS2^505R^-HS1^248R^* line, an A6-like segment is present in *HS2^505R^-SV40-HS1^248R^* males (Figure 5B, S4). However, blocking is not fully restored to the level of *HS1^248R^* alone. In *HS2^505R^-SV40-HS1^248R^* males, the A6 tergite is slightly reduced in size while the sternite is absent. In the embryonic CNS, Abd-B expression in PS11 is clearly elevated, but less so than either that observed in either *HS2^505R^-HS1^248R^* or *F7^attP50^* (Figure 5C). To confirm that the partial reactivation of blocking activity is due to a reduction in transcriptional readthrough, we generated a similar construct, *HS2^505R^-mSV40-HS1^248R^*, in which the polyadenylation sequences in the SV40 fragment were mutated. In contrast to *HS2^505R^-SV40-HS1^248R^*, the insulating activity of *HS2^505R^-mSV40-HS1^248R^* is substantially compromised and only a residual A6 tergite is observed (Figure 5B, S4).

In these experiments the SV40 termination element will not only suppress transcription through *HS1*, but also into the neighboring *iab-7* regulatory domain. To rule out possibility that the SV40 element “rescues” the blocking activity of *HS1* by reducing transcription into the *iab-7* regulatory domain, we placed it downstream of *HS1* instead between *HS2* and *HS1*. Fig. 5B and S4 shows that just like *HS2^505R^-HS1^248R^, HS2^505R^-HS1^248R^-SV40* has no boundary function: A6 is absent in males carrying the replacement, indicating that cells in PS11/A6 have assumed a PS12/A7 identity. The *Abd-B* expression pattern in the embryonic CNS also closely matches that of *HS2^505R^-HS1^248R^* line (Fig. 5C). Similar results were obtained when the same construct (*SV40-HS1^249^-HS2^505^*) was inserted in the forward orientation so that the *HS2* element is “pointing” towards *iab-6* (Fig 5, S4).

To provide further evidence that transcriptional read-through from *HS2* is responsible for disrupting the blocking activity of *HS1*, we generated a replacement, *HS2^505^-HS1^248R^*, in which the 5’→3’ orientation of *HS2* was inverted with respect to *HS1*. If transcription from *HS2* is unidirectional, then the blocking activity of *HS1* should be unaffected when *HS2* is “pointing” away from *HS1*. Consistent with this prediction, *HS1* retains blocking activity in *HS2^505^-HS1^248R^*. Fig. 5C shows that the pattern of Abd-B expression in the embryonic CNS in both PS10 and PS11 is reduced by *HS2^505^-HS1^248R^*, while the cuticle phenotype in adult males resembles *HS1^248^* (Figure 5B). Taken together, these results show that read-through transcription from *HS2* is responsible for the inactivation of *HS1* boundary function.

### Transcription from *HS2* promoter cannot overcome five dCTCF sites

Transcriptional read-through by RNA Polymerase II (Pol II) could disrupt *HS1* boundary activity by temporarily displacing dCTCF as well as other boundary associated factors. If this is the case, it seemed possible that the presence of additional dCTCF sites in *HS1* would counteract the effects of transcription from *HS2* either by helping to maintain boundary function as Pol II reads through or by inhibiting Pol II elongation. To test this possibility, we inserted four additional dCTCF sites in an *HS1* Sph I restriction site located close to the endogenous dCTCF site. Fig 5B shows that the addition of 4 dCTCF sites (*HS2^505R^-HS1^248R^-CTCF^x4^*) is sufficient to rescue the blocking activity of *HS1*. As observed for *HS1* alone, *HS2^505R^-HS1^248R^-CTCF^x4^* males have a tergite and sternite whose morphology resembles that seen in A5. Likewise, the pattern of *Abd-B* expression in the CNS in *HS2^505R^-HS1^248R^-CTCF^x4^* is similar to that observed for *HS^248^* (Fig. 5C).

### Segmental regulation of the *HS1* boundary by *HS2* dependent transcriptional read-through

Taken together, the findings in the previous sections suggest that *Fub-1* likely functions as a boundary in its endogenous context; however, its’ boundary activity is segmentally regulated by transcriptional read through from *HS2*. This model makes two predictions. First, the activity of the *HS2* promoter should be controlled by the *bxd/pbx* regulatory domain. Second, when transcriptional read-through is prevented, *Fub-1* blocking activity would be expected to interfere with *Ubx* regulation by the *bxd/pbx* regulatory domain. We have tested these predictions

#### HS2 promoter activity is controlled by the bxd/pbx regulatory domain

In order to test the first prediction, a probe that specifically recognizes transcripts originating from the *HS2* promoter is required. As noted above, Pease et al., 2013 found that there are (at least) two distinct “sense” lncRNAs, *bxd* and *F1^HS2^*, expressed under the control of the *bxd/pbx* regulatory domain that could potentially read-through *HS1* and be complementary to sequences in the region between *HS1* and the *Ubx* promoter. For this reason, transcripts originating from the *HS2* promoter and extending through *HS1* towards the *Ubx* promoter cannot be unambiguously identified as being derived from *HS2*.

On the other hand, antisense (proximal to distal) transcripts from this region are not detected in *WT* embryos. Thus, it would be possible to assay the promoter activity of *HS2* by reversing its orientation in the genome. For this purpose, we generated two different *Fub-1* replacements in which the 5’→3’ orientation of the *HS2* element was reversed so that it would generate “antisense” transcripts. In the first, we inserted a fragment, *HS1^248^-HS2^505R^*, into *F1^attP^* that contains both *Fub-1* hypersensitive sites (Fig.5D). However, the 5’→3’ orientation of *HS2* in this fragment was reversed so that it is pointing away from *HS1*. In this case transcripts expressed from the *HS2* promoter would extend into the *bxd/pbx* domain, instead of transcribing through *HS1*. In the second, we inserted a *Fub-1* fragment in *F1^attP^* containing both hypersensitive sites, *HS2^505R^-HS1^248R^*, but in the reverse orientation so that transcription from *HS2* would read-through *HS1* towards the *bxd/pbx* regulatory domain (Fig.5D).

Antisense transcripts in *WT, HS1^248^-HS2^505R^* and *HS2^505R^-HS1^248R^* embryos were then visualized using smFISH probes spanning a 1463 bp *bxd* sequence located just distal to the *att*P site (Fig.5D). Fig. 6 shows that antisense transcripts complementary to the smFISH probes are not detected in *WT* embryos. In contrast, segmentally restricted antisense transcripts are expressed in both of the *Fub-1* replacements. Like the *DsRed* and *GFP* reporters described above (Fig. 3), the antisense transcripts are detected in PS6/A1 through PS13/A8. These findings show that the *HS2* promoter in the *Fub-1* replacements is regulated by the *bxd/pbx* domain, while it is insulated from regulatory elements in the *abx/bx* domain. The transcripts levels in *HS1^248^-HS2^505R^* and *HS2^505R^-HS1^248R^* embryos are not, however, equivalent. The signal is noticeable greater in *HS1^248^-HS2^505R^* than it is in *HS2^505R^-HS1^248R^.* This could reflect the fact that the smFISH probes are complementary to sequences immediately adjacent to HS2 in the *HS1^248^-HS2^505R^* replacement, whereas the polymerase must traverse *HS1* in the *HS2^505R^-HS1^248R^* replacement. Alternatively, since *HS1* is located between the *HS2* promoter and the *bxd/pbx* regulatory domain, it may attenuate regulatory interactions, and at least the initial activation in PS6 may depend upon the enhancers thought to be located between the *Ubx* promoter and *Fub-1*.

**Figure 6.**
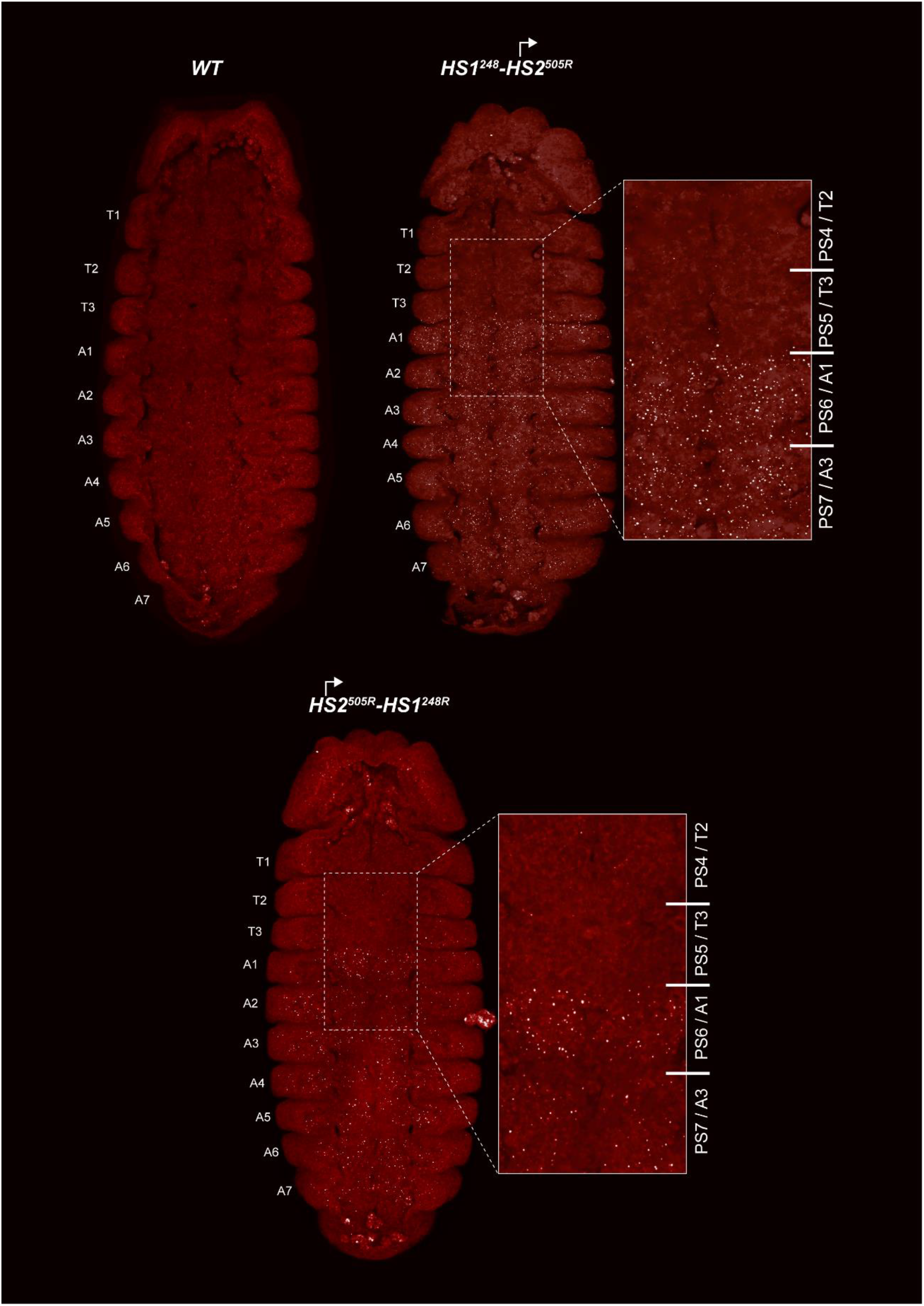
*HS2* drives transcription in tissue-specific manner. smFISH of stage 14 embryos of indicated genotype. In *WT* embryos no antisense (proximal to distal) transcripts were detected. By contrast, in *HS1^248^-HS2^505R^* and *HS2^505R^-HS1^248R^* antisense transcripts are detected from PS6/A1 through PS13/A8. Note that the intensity of smFISH signal is higher in *HS1^248^-HS2^505R^* than in *HS2^505R^-HS1^248R^*.

#### Transcriptional read through is required to abrogate HS1 blocking activity in PS5/A1

The second prediction is that transcriptional read-through from *HS2* into *HS1* is required to relieve the blocking activity of *HS1*, enabling regulatory interactions between the *bxd/pbx* domain and *Ubx* in PS5/A1. This prediction also holds. As shown in Fig. 2, the morphology of the A1 segment in *HS2^505R^-HS1^248R^* replacement flies is similar to that in *WT*. In this replacement transcripts originating in *HS2* read through *HS1* into the *bxd/pbx* regulatory domain. A different result is obtained for the replacement, *HS1^248^-HS2^505R^*, in which the *HS2* promoter is directed away from HS1. *HS1^248^-HS2^505R^* flies show evidence of a strong loss-of-function (LOF) transformation (Fig.5E). This replacement transforms the first abdominal segment into the third thoracic: the first abdominal tergite is reduced or absent, in addition, the posterior third thoracic segment is partly transformed to a posterior second thoracic segment – the halters enlarged and pointing downward (Fig. 2B). These phenotypes are known as *bithoraxoid* (*bxd*) and *postbithorax* (*pbx*) respectively (Bender et al., 1983). *HS1^248^-HS2^505R^* homozygotes also display very low viability and sterility.

## DISCUSSION

While a very large percentage of the Pol II transcripts in multicellular eukaryotes are lncRNAs, what roles they play in the expression coding mRNAs and in development largely unknown. Here we have investigated the functional properties of a lncRNA gene, *F1^HS2^*, that is located in the *bxd/pbx* regulatory domain of *Drosophila* BX-C. As has been observed for many other lncRNA genes, the RNA sequences encoded by *F1^HS2^* are not well conserved and it is the functional properties of the two *F1^HS2^* regulatory elements, *HS1* and *HS2*, that are important. We show here that *HS1* is a dCTCF dependent boundary element, while *HS2* is a developmentally regulated promoter that directs transcription through *HS1* inactivating its boundary function in parasegments that require *Ubx* function.

The arrangement of the three homeotic genes and the nine parasegment specific regulatory domains in the *Drosophila* BX-C complicates the coordination between the 3D organization of the complex and the requirements for regulatory interactions. In order to specify segment identity, the nine regulatory domains are segregated into topologically independent loops by boundary elements. This physical organization helps to block crosstalk between initiation elements in adjacent domains in the early embryo, while later in development it helps restrict the spread of PcG silencing from inactive to active domains (Kyrchanova et al., 2015; Maeda and Karch, 2015). However, in order to direct the proper expression of their target homeotic gene, all but three of the regulatory domains must be able to bypass one or more intervening boundary elements. In the case of the *Ubx* gene, the *bxd/pbx* regulatory domain is separated from its target promoter by the *Fub-1* boundary, and by a second boundary element located close to the *Ubx* promoter.

The *Fub-1* boundary is subdivided into two elements, *HS1* and *HS2*. *HS1* like other boundaries in BX-C has a dCTCF site, is bound by dCTCF *in vivo* and is able to function as a generic insulator. While *HS2* may contribute to the boundary function of *Fub-1*, it does not have insulating activity on its own. Instead, it has promoter activity, and can function as an enhancer trap. In its normal context, the *HS2* promoter is controlled by the *bxd/pbx* regulatory domain. The *bxd/pbx* regulatory domain is set in the *off* state in parasegments (segments) anterior to PS6/A1 at the blastoderm stage and is repressed during the remainder of development by PcG dependent silencing. In PS6/A1 (and more posterior segments) the *bxd/pbx* regulatory domain is set in the *on* state, activating the stage and tissue specific enhancers in this domain. These stage and tissue specific enhancers activate the *HS2* promoter in PS6/A1 and more posterior parasegments and transcription from this promoter through *HS1* disrupts *Fub-1* boundary function (Fig. 7). This enables stage and tissue specific enhancers in *bxd/pbx* to activate *Ubx* expression in PS5/A1 and more posterior parasegments. In addition to the *F1^HS2^* lncRNA, at least one other segmentally regulated lncRNA, *bxd* lncRNA, is transcribed through the *Fub-1* boundary and thus could also disrupt the blocking activity of the *Fub-1* boundary in PS6/A1 and more posterior parasegments.

**Figure 7.**
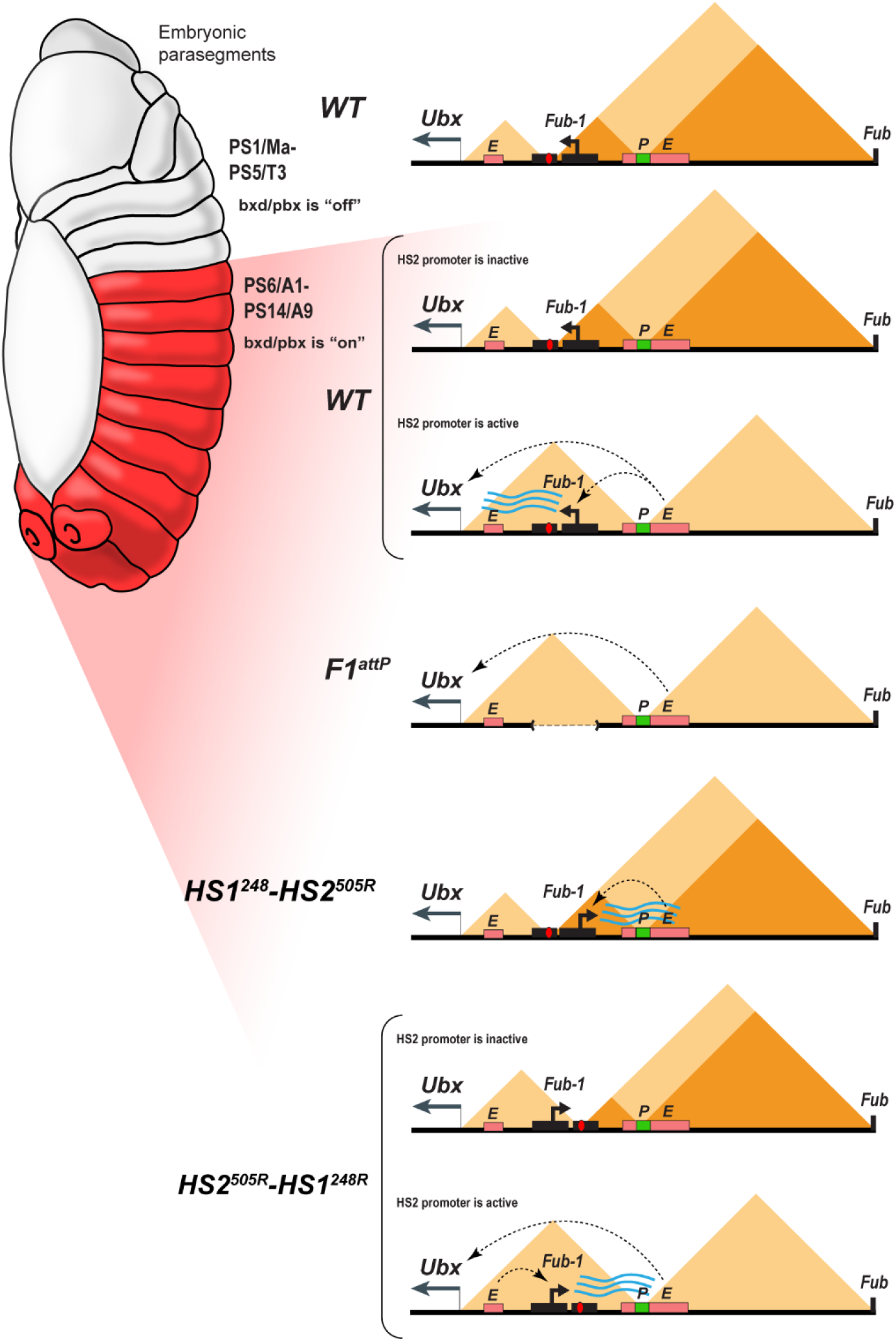
Model showing changes in TAD organization. <u>Top</u>: In *WT* the 3D organization of the *bxd/pbx* regulatory domain transitions from one state to another when it activates *Ubx* expression (see also (Mateo et al., 2019)). In PS5 (T3) and more anterior parasegments, the *bxd/pbx* domain is encompassed in a large TAD that extends from *Fub-1* to *Fub* (large triangle). Within this large TAD there are two HDIC domains. One extends from *Fub-1* to the *bxd* PRE, while the other extends from the *bxd* PRE to *Fub.* In PS6 and more posterior parasegments the 3D organization of the domain is dynamic and depends upon whether the *Fub-1 HS2* promotor is active or not. When the promoter is inactive (in between bursts), the *Fub-1* boundary is expected to be functional and that TAD/HDIC organization would resemble that in more anterior parasgments. When the enhancers in *bxd/pbx* activate *HS2*, transcriptional readthrough disrupts *Fub-1* boundary activity generating a new TAD organization that links enhancers in *bxd/pbx* to the *Ubx* promoter. <u>*F1^attP^*</u>: Deletion of the *Fub-1* boundary eliminates the large *Fub-1←→Fub* TAD in all parasegments. In this case, an element upstream of the *Ubx* promoter functions to insulate *abx/bx* from *bxd/pbx* in PS5/T3. <u>*HS1^248^-HS2^540R^*</u>: In this replacement *HS2* is inverted so that the *HS2* promoter is pointed away from *HS1* towards the *bxd/pbx* regulatory domain. Though *bxd/pbx* activates the promoter in PS6/A1 and more posterior parasegments, *Fub-1* boundary activity is not disrupted. <u>*HS2^540R^-HS1^248R^*</u>:_In this replacement, *Fub-1* is inverted. Though *HS2* promoter activity in PS6 (A1) and more posterior parasegments is reduced in this configuration compared to WT, it is sufficient to disrupt *Fub-1* boundary activity and enable *bxd/pbx* to regulate *Ubx* expression.

Several lines of evidence are consistent with this mechanism. When *HS1* is used to replace the *Fab-7* boundary, it functions like other generic fly boundaries and blocks crosstalk between the *iab-6* and *iab-7* initiators, but does not support boundary bypass. This blocking activity is also orientation independent. In contrast, neither *HS2* alone nor the intact *Fub-1* element are able to block crosstalk between *iab-6* and *iab-7* and both replacements exhibit the same GOF transformations as the starting *F7^attP^* replacement platform. In these *Fab-7* replacement experiments the defects in *Fub-1* blocking activity arise from transcriptional readthrough from the *HS2* promoter. This conclusion is supported by two lines of evidence. First, introducing the SV40 transcription termination polyadenylation element in between *HS2* and *HS1* partially rescues the blocking defect. Rescuing activity is not due to the increased distance between *HS2* and *HS1* as a SV40 element with mutations in the polyadenylation signal significantly compromising its rescuing activity. In addition, the SV40 termination element must be located between *HS1* and *HS2* as rescue is not observed when it is placed downstream of *HS1*. Second, when *HS2* is inverted so that transcription proceeds away from *HS1*, boundary function is also restored.

A similar mechanism appears to regulate *Fub-1* boundary activity in its endogenous context; however, in this case boundary activity is segmentally regulated. In PS5 and more anterior parasegments, the *bxd/pbx* regulatory domain is maintained in an “*off”* state by a PcG based mechanism. As a consequence, the *Fub-1 HS2* promoter is inactive (Fig.7). In PS6 (A1) (and more posterior parasgements) the *bxd/pbx* domain is in the “*on”* state and it activates transcription from the *Fub-1 HS2* promoter. When *HS2* is oriented so that transcription proceeds through *HS1*, boundary activity is abrogated in PS6/A1 and more anterior parasegments/segments (Fig. 7). This enables enhancers in the *bxd/pbx* regulatory domain to activate *Ubx* transcription. Consistent with parasegment specific differences in the 3D topology imaging studies of fly embryos by Mateo et al., showed that the *Ubx* region of BX-C undergoes a reorganization in PS5/T3 and PS6/A1 (Mateo et al., 2019). This rearrangement would be blocked when *HS2* is oriented away from *HS1*. In this case, boundary activity is retained and the stage and tissue enhancers in the *bxd/pbx* regulatory domain are unable to activate *Ubx* expression (Fig. 7). This leads to a LOF phenotype much like that observed in PS11/A6 in *Fab-7* replacement experiments when the *HS1* boundary blocks crosstalk but does not support bypass.

While it is clear that *Fub-1* boundary function is subject to negative and positive regulation by elements in the *bxd/pbx* domain, and that this is important for the proper specification of PS6/A1, there are a number of outstanding questions. One is whether transcription from the *HS2* promoter is necessary to inactivate *HS1* earlier in development. As described above, *bxd* lncRNA transcripts emanating from a promoter in *pbx* extend through both *HS2* and *HS1*. If *HS1* is sensitive to read-through from *HS2*, one would think that it would also be disrupted by transcriptional read-through of the *bxd* lncRNA. However, since Pease et al found that transcription of the *bxd* lncRNA is not required for normal development, it would appear that this lncRNA is dispensable, at least when *Fub-1 HS2* is present. While we do not know the pattern of *F1^HS2^* or *bxd* lncRNAs expression after embryogenesis, the fact that reversing *HS2* results in morphological defects in adults would argue that *F1^HS2^* is likely expressed during the larval and pupal stages.

Another question is raised by the Micro-C contact pattern in *WT* and the *Fub-1* deletion. In *WT*, *Fub-1* and *Fub* define a TAD that encompasses *bxd/pbx*. *Fub-1* also corresponds to a boundary for two smaller HDIC domains, one with a second endpoint close to the *Ubx* promoter and another with a second endpoint close to the *bxd* PRE. In the *Fub-1* deletion, the proximal endpoint of the *bxd/pbx* TAD shifts to the *Ubx* promoter region, while the two HDIC domains collapse into one. If the function of *HS2* is to inactivate *HS1* in the posterior 2/3rds of the embryo, one would expect that this would disrupt the *Fub-1 ←→ Fub* TAD, much like the *Fub-1* deletion. There could be several reasons for this discrepancy. Since the *HS2* promoter should not be active in cells anterior to PS6/A1, it is possible that these cells are responsible for generating the *Fub-1 ←→ Fub* TAD seen in the Micro-C experiments. Alternatively, the disruption in *HS1* function might only occur during a burst of transcription (Fig. 7). Recent studies have shown that transcription of most genes is not continuous but rather occurs in discrete bursts (Rodriguez and Larson, 2020; Tunnacliffe and Chubb, 2020). The frequency, duration and amplitude of the burst can vary substantially. In some experimental systems, bursts can last several minutes or longer while the intervals between bursts may be only 10s of seconds (Bothma et al., 2014; Fukaya et al., 2016). For other genes, the intervals between bursts can be 30 min or more and the bursts may last only a minute or two (Alexander et al., 2019). If disruption of *HS1* blocking activity is directly coupled to the act of transcription (as appears to be the case0, then there may only be small windows of time in which the activity of the *Fub-1* boundary is compromised and *bxd/pbx* enhancers can activate the *Ubx* promoter. In this case, the *Fub-1* boundary would flip back and forth from an inactive and active state (Fig.7).

A related issue would be the mechanism activating *HS2* transcription when the *Fub-1* boundary is reversed (the *HS1^248^-HS2^505R^* replacement) so that *HS1* is interposed between *HS2* and the *bxd/pbx* domain. One would have expected that *HS1* would block the *bxd/pbx* enhancers from activating *HS2* transcription. In this case, boundary function would be retained, resulting in a loss-of-functtion transformation of PS6/A1 into PS5/T3 because in would prevent *bxd/pbx* from regulating *Ubx*. One possibility is that the putative enhancers in the region between the *Ubx* promoter and *Fub-1* (see *2218S e*nhancer in Sup Fig. 2) are able to activate a sufficient level of *HS2* dependent transcription to disrupt HS1 boundary function. An alternative/additional possibility is that the blocking activity of *HS1* is not sufficient to completely suppress regulatory interactions between *bxd/pbx* and *HS2.* Potentially consistent with this idea is the finding that the orientation of the *Fub-1* element in the *Fab-7* replacements does not seem to be important. When *HS2* is adjacent to *iab-6* it inactivates the blocking activity of *HS1*. This would be the expected result since reporters in BX-C are activated when the regulatory domain in which they are inserted is turned *on* in the early embryo. In this case, *iab-6* would drive *HS2* expression in PS11(A7) cells and this would inactivate the blocking activity of *HS1* in cell in this parasegment, giving a GOF transformation of PS11/A6. The unexpected result is that a GOF transformation of PS11/A6 into PS12/A7 is also observed when the *Fub-1* replacement is inserted so that *HS2* is next to *iab-7*. Since *iab-7* is *“off”* in PS11, this would imply that *HS2* must be activated by the *iab-6* regulatory domain. A similar scenario could be at play when *Fub-1* is inserted in the opposite orientation in its endogenous location: *HS1* blocks but does not completely eliminate interactions between the *bxd/pbx* enhancers and the *HS2* promoter. This is probably not unexpected. While interposed boundaries significantly reduce enhancer:promoter contacts, they are not completely eliminated (Fukaya et al., 2016). In addition, once activated, transcription from *HS2* through *HS1* should weaken *HS1* boundary activity and this in turn would facilitate further transcription.

Perhaps the most puzzling question is what function does *Fub-1* play in BX-C regulation? Since there is no obvious GOF phenotype when *Fub-1* is deleted, it differs from other boundaries in BX-C that have been analyzed in that would appear to be dispensable. On the other hand, when present, it must be inactivated in PS6/A1 (and more posterior parasegments) so that the *bxd/pbx* domain can regulate *Ubx* expression. Interestingly, like *cis*-acting elements for many other lncRNAs, both of the *Fub-1* boundary sub-elements are conserved in the *Sophorphora* sub-genus (Sup Fig.3). It is also conserved in *D.virilis* which is a member of the *Drosophila* sub-genus. *D.virilis* differs from *melanogaster* in that the *Ubx* gene and its two regulatory domains, *abx/bx* and *bxd/pbx*, are separated from the *abd-A* and *Abd-B* genes. In spite of this rearrangement, the *Fub-1* sequences are conserved. The fact that *Fub-1* is present in distantly related species would argue that *Fub-1* has an important function under some type of selective condition (that we have yet to discover) which is commonly encountered by a wide range of *Drosophila* species.

## MATERIALS AND METHODS

### Key resources table

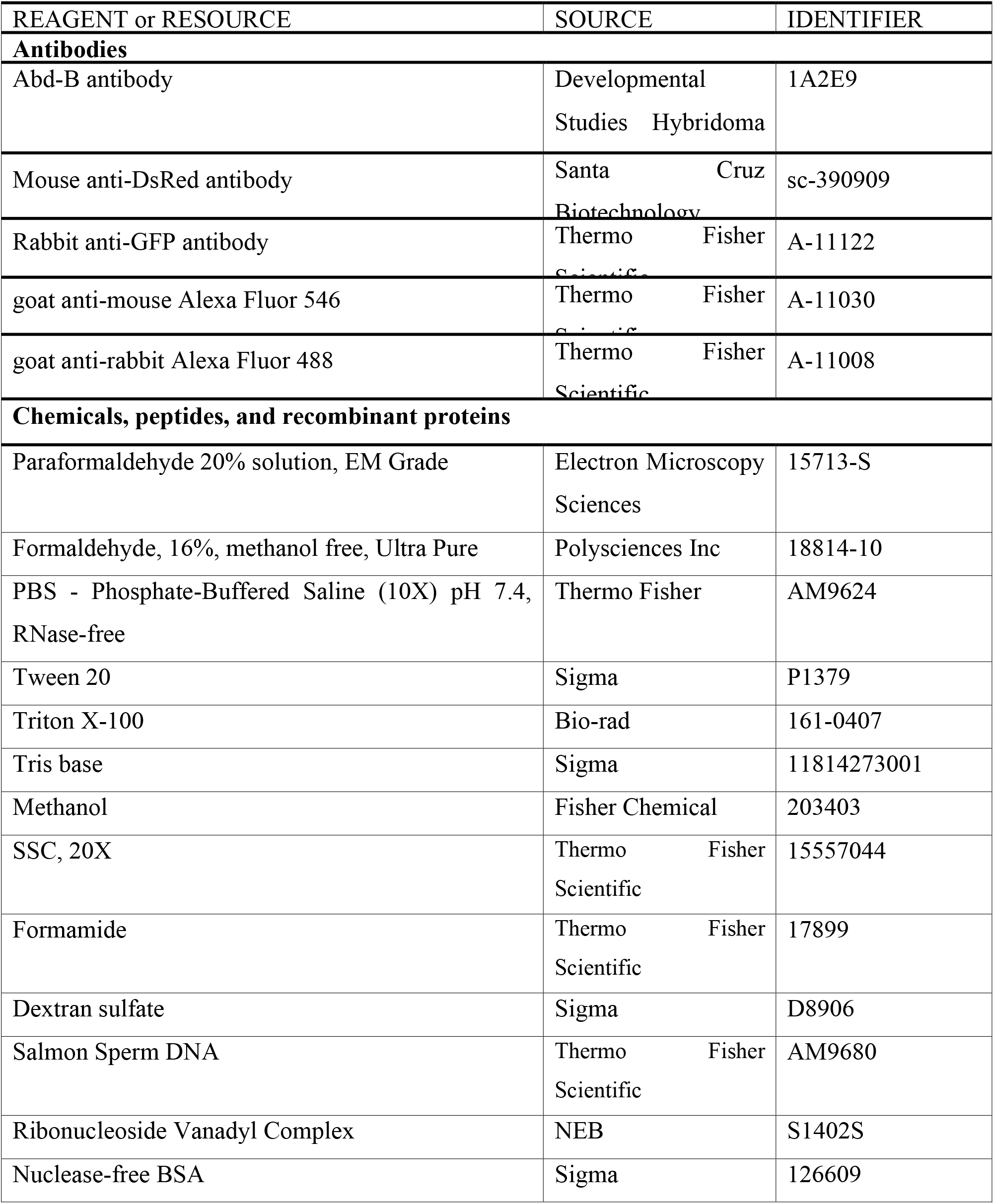
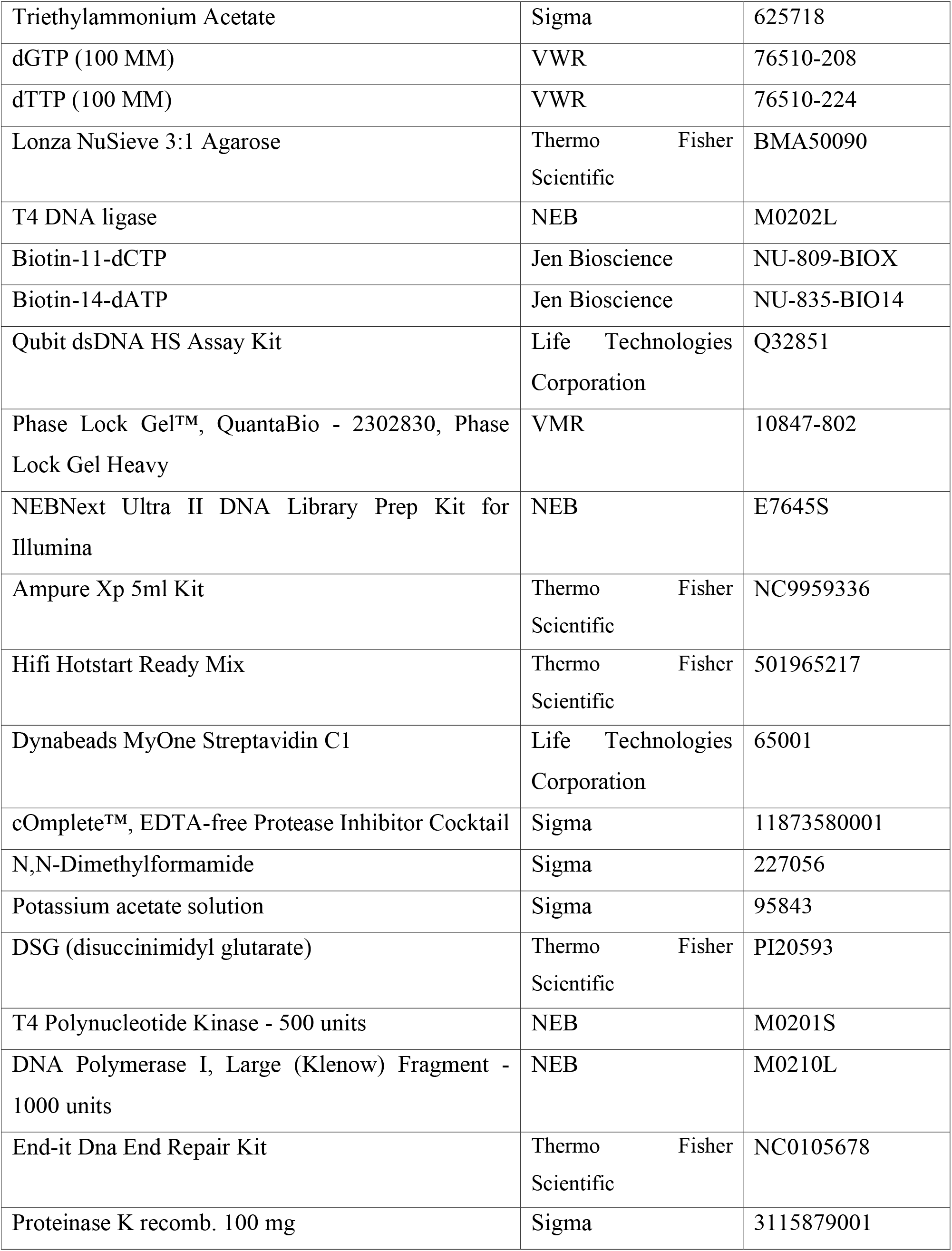
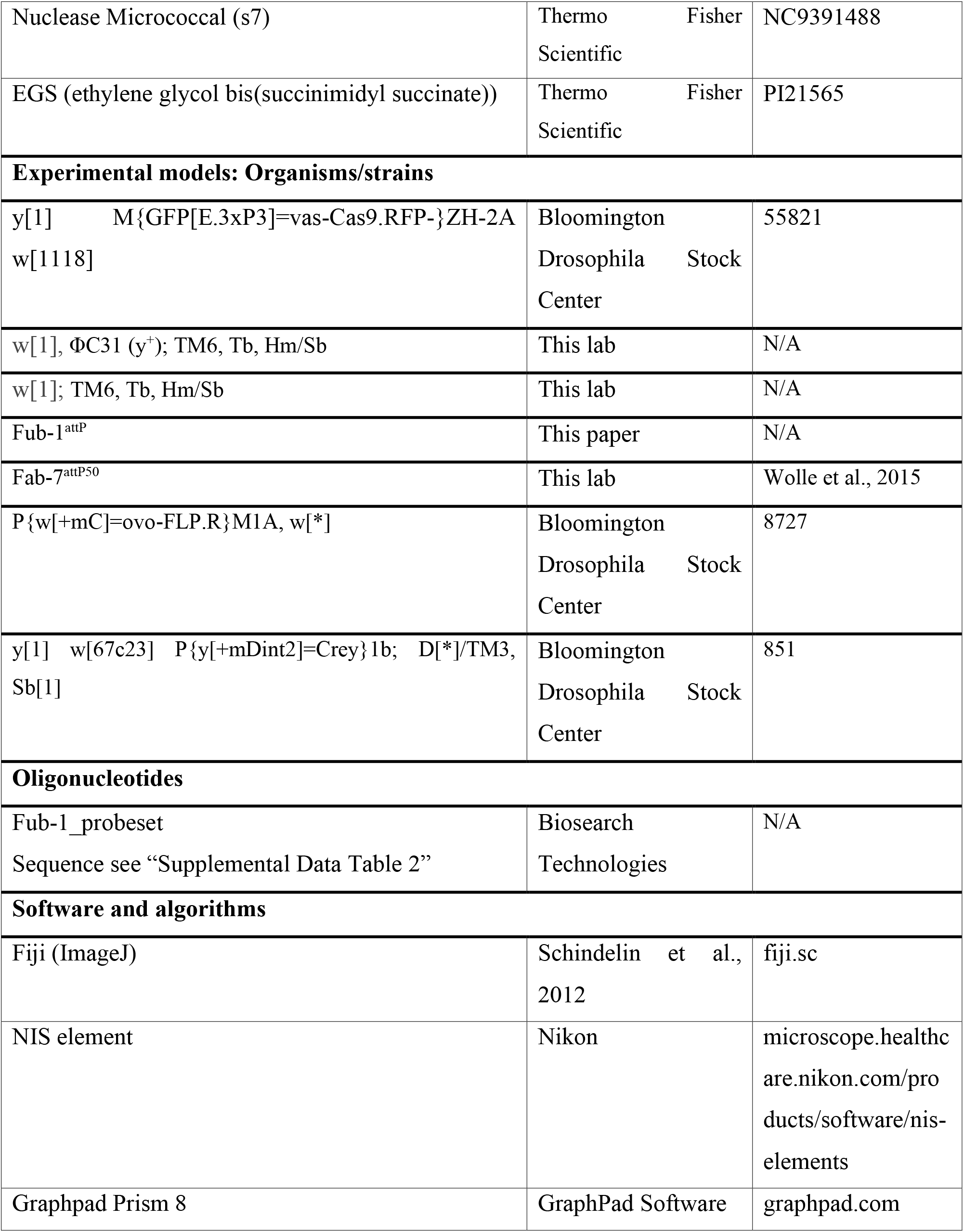
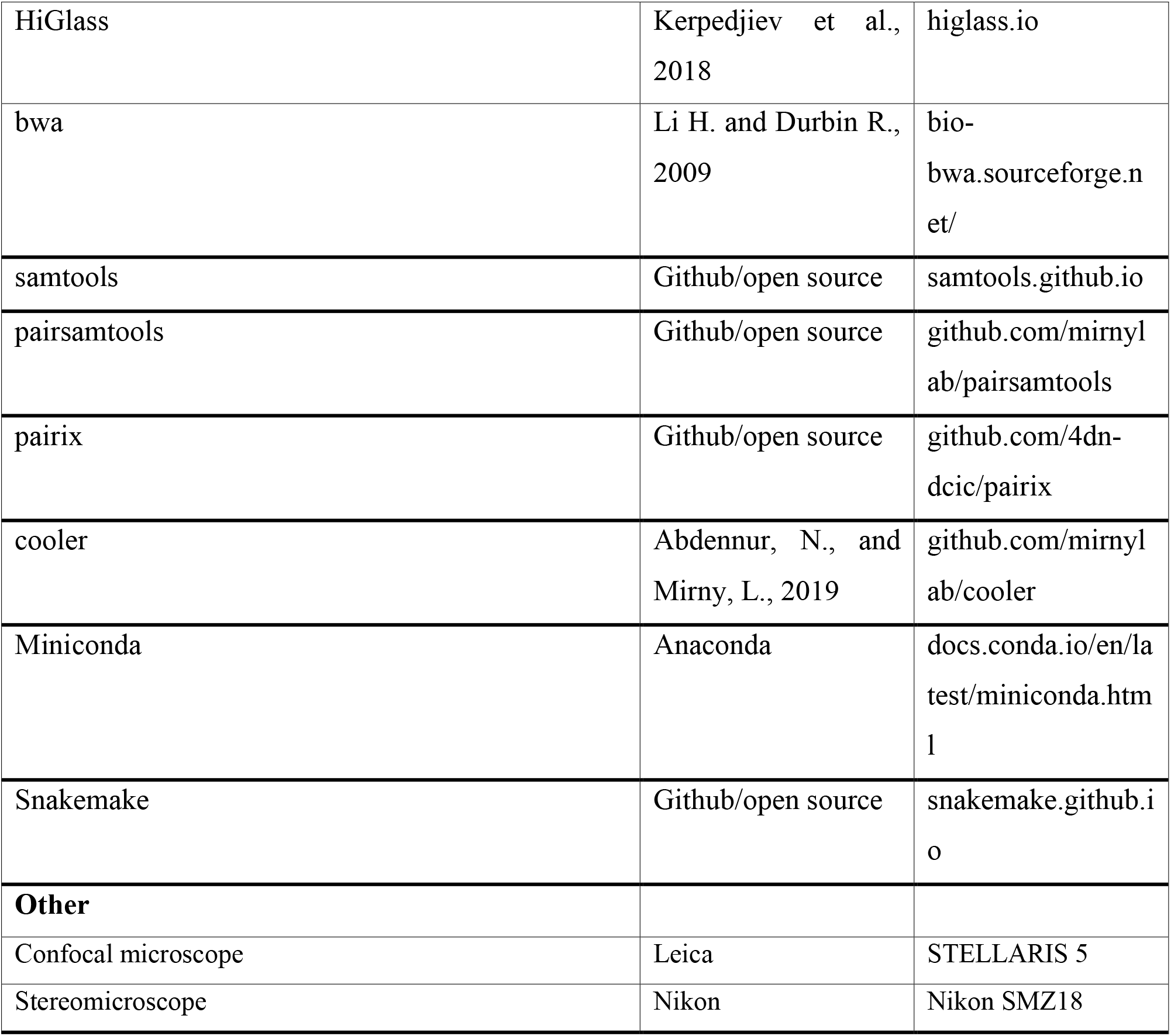

### Fly strains

The following stocks were used in these experiments: y[1] M{GFP[E.3xP3]=vas-Cas9.RFP-}ZH-2A w[1118] (BDSC_55821), w[1], ΦC31 (y^+^); TM6, Tb, Hm/Sb, w[1]; TM6, Tb, Hm/Sb, P{w[+mC]=ovo-FLP.R}M1A, w[*] (BDSC_8727), y[1] w[67c23] P{y[+mDint2]=Crey}1b; D[*]/TM3, Sb[1] (BDSC_851) was used for all transgenesis. y[1] M{GFP[E.3xP3]=vas-Cas9.RFP-}ZH-2A w[1118] (BDSC_55821) was used for CRISPR/Cas9 genome editing. y[1] w[67c23] P{y[+mDint2]=Crey}1b; D[*]/TM3, Sb[1] (BDSC_851) was used for Cre/loxP recombination to remove DsRed from CRISPR-generated DsRed insertions. w[1]; TM6, Tb, Hm/Sb, P{w[+mC]=ovo-FLP.R}M1A, w[*] (BDSC_8727) was used for Flp/frt recombination to remove GFP from ΦC31-mediated insertions. All flies were kept on standard fly cornmeal-molasses-yeast media.

### Generation of *F1^attP^* by CRISPR-Cas9–induced homologous recombination

For generating double-stranded DNA donor template for homology-directed repair, we used pHD-DsRed vector (Addgene plasmid no. 51434). The final plasmid contains genetic elements in the following order: [bxd proximal arm]-[attP]-[loxP]-[3×P3-dsRed-SV40polyA]-[loxP]-[bxd distal arm] (fig.S1). Homology arms were PCR amplified from the Bloomington Drosophila Stock Center line no. 55821 genomic DNA using the following primers: AGCTCTCGAGAGCGATGGAACCGTTTCTG and GTAGATCTCGGTTTACGATCGACTGGC for the proximal arm (1001-bp fragment), and TTGCGGCCGCATTTCTACGTTTATAAGCTGTTAATC and AAGCATGCCCACAATGGAGGCTGC for the distal arm (1002-bp fragment). Targets for Cas9 were selected using “CRISPR optimal target finder”—the program from O’Connor-Giles Laboratory. The following gRNA sequences: AGTCGATCGTAAACCGGAG and TTATAGACGTAGAAATGTA, were inserted into the pU6-BbsI-chiRNA vector (Addgene plasmid no. 45946). A mixture of the donor vector (500ng/ml) and two gRNA vectors (100ng/ml each) was injected into embryos of y[1] M{GFP[E.3xP3]=vas-Cas9.RFP-}ZH-2A w[1118] (no. 55821 from the Bloomington Drosophila Stock Center). Injectees were grown to adulthood and crossed with w[1]; TM6/Sb line. Flies positive for dsRed signal in eyes were selected into a new separate line. The successful integration of the recombination plasmid was verified by PCR and corresponded to the removal of 1168 bp within the *Fub-1* region (genome release R6.22: 3R:16,748,143..16,749,310).

### Generation the replacement lines

For the *F1^attP^* replacement, the recombination plasmid was designed *de novo* and contains several genetic elements in the following order: [loxP]-[pl]-[frt]-[3P3-GFP]-[frt]-[attb] (fig. S1). All elements were assembled within the pBluescript SK vector. loxP site is located before polylinker [pl] and in combination with the second site, which is located in the platform, used for excision of *DsRed* marker gene and plasmid body. Two frt sites are used to excise GFP marker. DNA fragments used for the replacement experiments were generated by PCR amplification and verified by sequencing (presented in the Supplementary Methods).

The strategy of the Fab-7 replacement lines creation is described in detail in (Wolle et al., 2015).

### Cuticle samples preparation

3 day adult male flies were collected in 1.5ml tubes and stored in 70% ethanol at least 1 day. Then ethanol was replaced with 10% KOH and flies were heated at 70°C for 1h. After heating flies were washed with dH2O two times and heated again in dH2O for 1h. After that digested flies were washed three times with 70% ethanol and stored in 70% ethanol. The abdomen cuticles were cut off from the rest of the fly using fine tweezer and a needle of an insulin syringe and put in a droplet of glycerol on a microscope slide. Then the abdomens were cut longitudinally on the mid-dorsal side through all of the tergites with the syringe. Then the cuticles were flattened with a coverslip. Photographs in the bright or dark field were taken on the Nikon SMZ18 stereomicroscope using Nikon DS-Ri2 digital camera, processed with Fiji bundle 2.0.0-rc-46.

### Embryo Immunostaining

Embryo Immunostaining was performed as previously described (Deshpande et al., 1999). Primary antibodies were mouse monoclonal anti-Abd-B at 1:40 dilution (1A2E9, generated by S. Celniker, deposited to the Developmental Studies Hybridoma Bank), mouse monoclonal anti-DsRed at 1:50 dilution (Santa Cruz Biotechnology) and rabbit polyclonal anti-GFP at 1:500 dilution (Thermo Fisher Scientific). Secondary antibodies were goat anti-mouse Alexa Fluor 546 (Thermo Fisher Scientific) and goat anti-rabbit Alexa Fluor 488 (Thermo Fisher Scientific). Stained embryos were mounted with VECTASHIELD antifade mounting medium (Vector Laboratories). Images were acquired on a Leica STELLARIS 5 confocal microscope and processed using ImageJ 1.53f51 (Schindelin et al., 2012).

#### Electrophoretic mobility shift assay

Electrophoretic mobility shift assay was performed using γ-32P–labeled DNA probes as described previously (Kyrchanova et al., 2018; Wolle et al., 2015).

### smFISH

smFISH was performed as previously described (Little et al., 2013). Quasar 670-conjugated *bxd* probes were ordered from LGC Biosearch Technologies. Sequences are listed in Supplementary Table 4). Embryos were collected overnight and then dechorionated in Clorox, fixed in 4% paraformaldehyde, and devitellinized in methanol. The hybridization with the FISH probes was performed overnight at 37°C in hybridization buffer (10% dextran sulfate, 0.01% salmon sperm single-strand DNA, 1% vanadyl ribonucleoside, 0.2% bovine serum albumin, 4×SSC, 0.1% Tween-20, and 35% formamide). The resulting preparations were washed twice with wash buffer, 1 h each, at 37°C and mounted in VECTASHIELD antifade mounting medium. Images were acquired on a Leica STELLARIS 5 confocal microscope with 40x HC PL APO CS2 1.3 NA oil immersion objective. Image denoising was performed using NIS elements “Advanced denoising” tool.

### Micro-C library construction

Embryos were collected on yeasted apple juice plates in population cages. Plates were laid for 4 hours, incubated for 12 hours at 25°C, then subjected to fixation. Embryos were collected in nylon mesh sieves, dechorionated for 2 mins in 3% sodium hypochlorite, rinsed with deionized water, and transferred to glass vials containing 5 mL PBST (0.1% Triton-X in PBS), 7.5 mL N-heptane, and 1.5mL fresh 16% formaldehyde. Crosslinking was carried out at room temperature for exactly 15 mins on an orbital shaker at 250rpm, followed by addition of 3.7 mL 2M Tris-HCl pH7.5 and shaking for 5 mins to quench the reaction. Embryos were washed twice with 15 mL PBST and subjected to secondary crosslinking. Secondary crosslinking was done in 10mL of freshly prepared 3mM final DSG and ESG in PBST for 45 mins at room temperature with passive mixing. Reaction was quenched by addition of 3.7mL of 2M Tris-HCl pH7.5 for 5 mins, washed twice with PBST. Embryos were snap-frozen and stored at −80°C until library construction.

Micro-C libraries were prepared as previously described (Batut et al., 2022) with the following modification: we used 50uL of 12-16hr embryos, non-sorted for each biological replicate. 60U of MNase was used for each reaction to digest chromatin to 80% mononucleosome vs 20% dinucleosome ratio. Libraries were barcoded, pooled and subjected to paired-end sequencing on an Illumina Novaseq S1 100 nt Flowcell (read length 50 bases per mate, 6-base index read).

### Micro-C data processing

Micro-C data for *D. melanogaster* were aligned to a custom genome edited from the Berkeley Drosophila Genome Project (BDGP) Release 6 reference assembly (dos Santos et al., 2015) with BWA-MEM (Li and Durbin, 2009) using parameters -S -P -5 -M. Briefly, the custom genome was designed to remove the visually obstructive ~5.6kb DIVER_I-int repeat element between Fub-1 and Fub. The specific region deleted was the entire repeat element, chr3R: 16756620-16762262 from the dm6 reference FASTA file. The resultant BAM files were parsed, sorted, de-duplicated, filtered, and split with pairtools (https://github.com/mirnylab/pairtools). We removed pairs where only half of the pair could be mapped, or where the MAPQ score was less than three. The resultant files were indexed with pairix (https://github.com/4dn-dcic/pairix). The files from replicates were merged with pairtools before generating 100bp contact matrices with Cooler (Abdennur and Mirny, 2020). Finally, balancing and mcool file generation was performed with Cooler’s zoomify tool. The mcool files were visualized using HiGlass (Kerpedjiev et al., 2018).

## ACKNOWLEDGMENTS

We are grateful to the Center for Precision Genome Editing and Genetic Technologies for Biomedicine of IGB RAS and the Core Facilities Center of IGB RAS for providing research equipment. We thank Tatyana Smirnova for technical assistance and advice on smFISH data acquisition and analysis. We thank Girish Deshpande, Tsutomu Aoki, Olga Kyrchanova and Oleg Bylino for insightful discussions throughout the course of this work. This work was supported by R35 GM126975 to P.S. and R01 GM118147 to M.L. Immunostaining analysis was supported by Russian Science Foundation, Project No. 19-74-30026 (to P.G.). Part of this work (bioinformatics analysis) was supported by Russian Science Foundation, Project No. 20-14-00201 (to Y.S.).

## AUTHOR CONTRIBUTION

Conceptualization, A.I., P.S. and P.G.; Methodology, A.I., P.S. and P.G.; Investigation, A.I. and X.B.; Writing – Original Draft, A.I. and P.S.; Writing – Review & Editing, A.I., P.S. P.G. and M.L; Funding Acquisition, P.S., P.G., Y.S. and M.L.; Resources, P.S. and P.G.; Supervision, P.S., P.G and M.L.

## COMPETING INTERESTS

The authors declare no competing interests.

## SUPPLEMENTARY MATERIALS

**Table S1.**
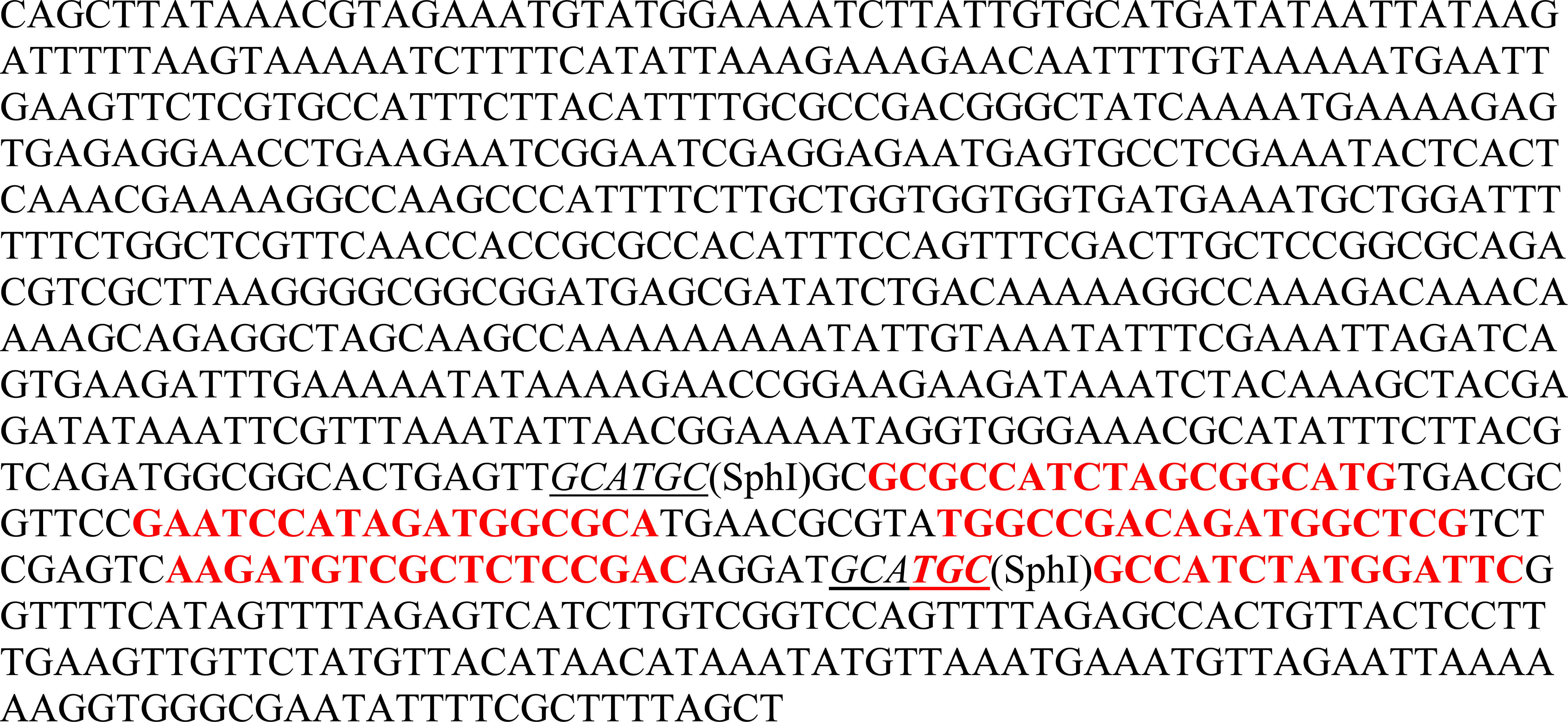
The sequence of *HS2^505R^-HS1^248R^-CTCF^x4^* fragment containing multimerized binding sites for dCTCF (CTCF×4). The binding sites are marked in red.

**Table S2.**
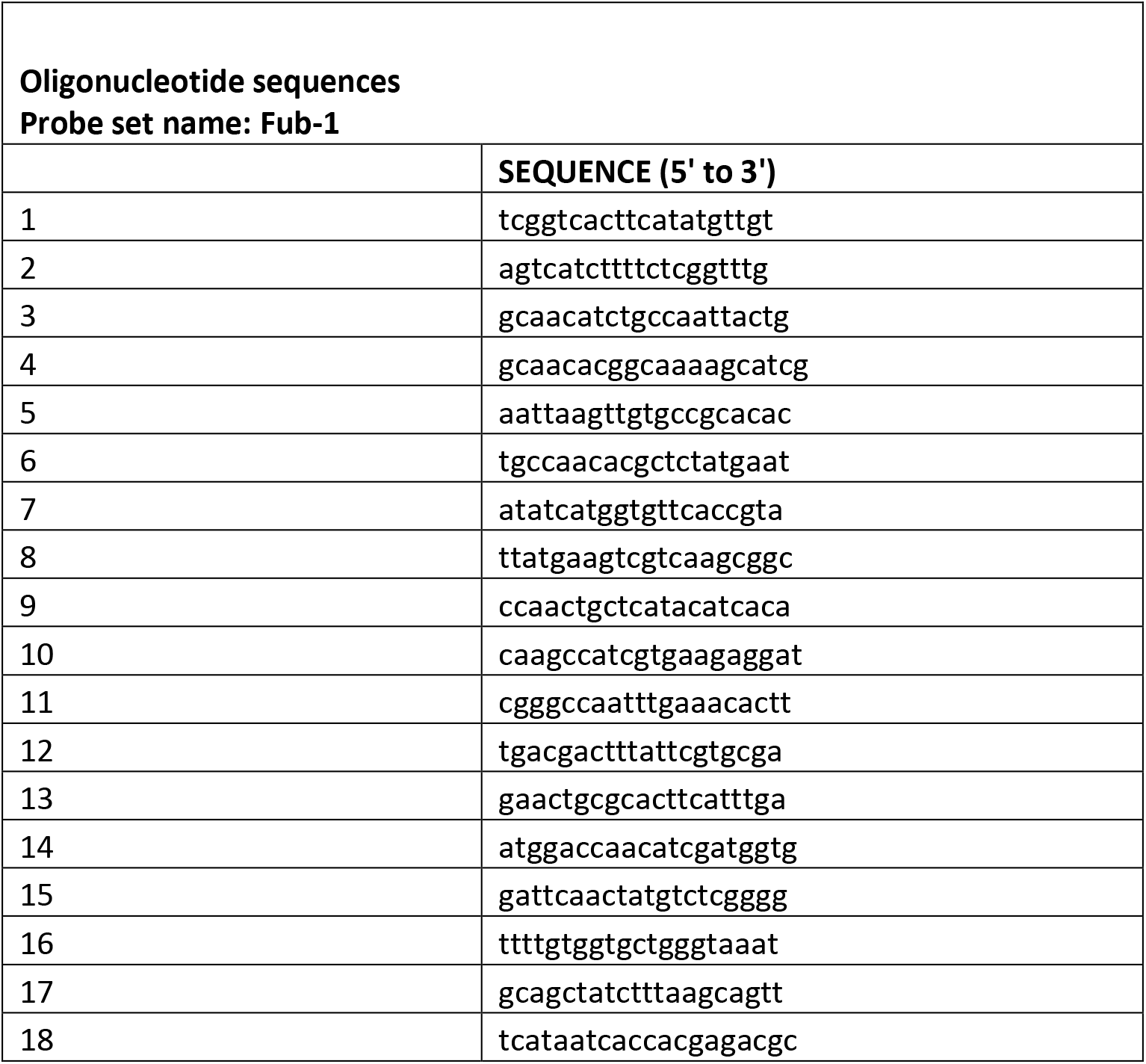
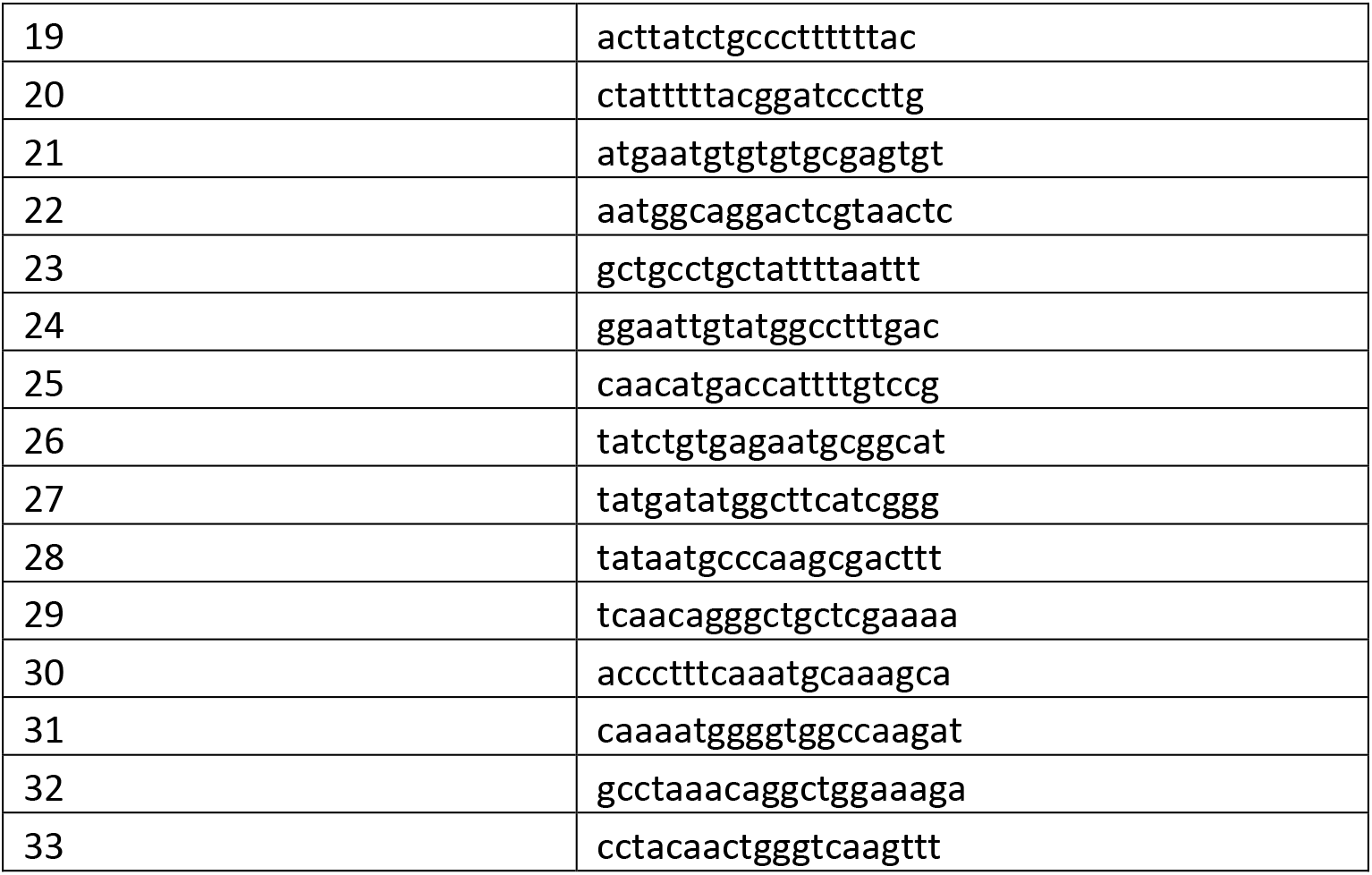
Sequences of smFISH probes covering 1463-bp region in *bxd* domain.

**Figure S1.**
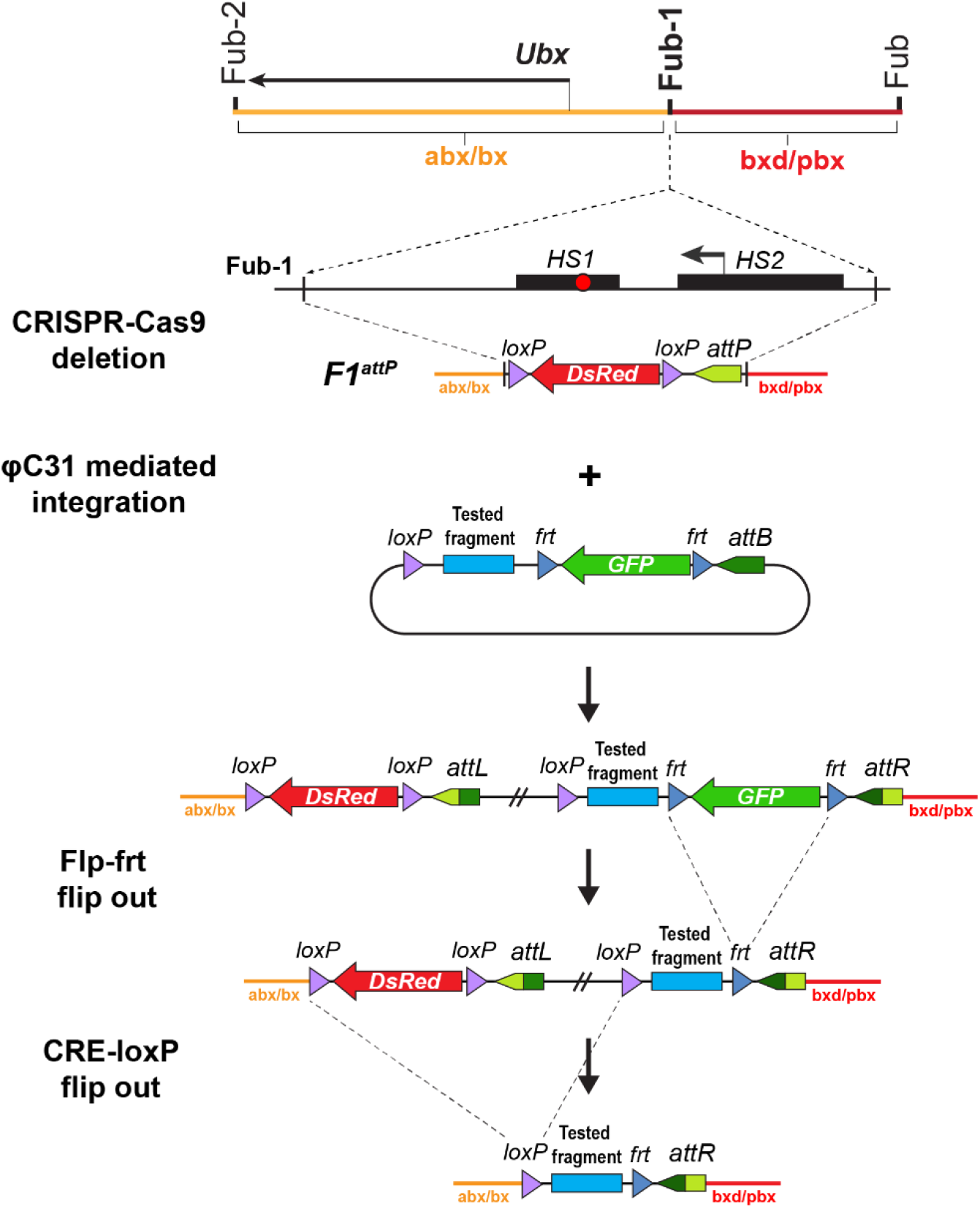
Strategy for creating *Fub-1* deletion and replacement lines. Top: schematic representation of the regulatory region containing the *Ubx* gene. The *F1^attP^* line was obtained through the substitution of a 1168-bp region with *attP* site and the *dsRed* gene, flanked by *loxP* sites. The coordinates of the deletion are dm6 3R:16,748,143..16,749,310. The plasmid that contains tested fragment and *GFP* marker gene was injected into the *F1^attP^* line. In the next step, *GFP* gene was excised by CRE-mediated recombination between the *frt* sites. During the final step, the *dsRed* gene was excised by recombination between the *loxP* sites. As a result, the tested elements were inserted in place of the 1168-bp deletion.

**Figure S2.**
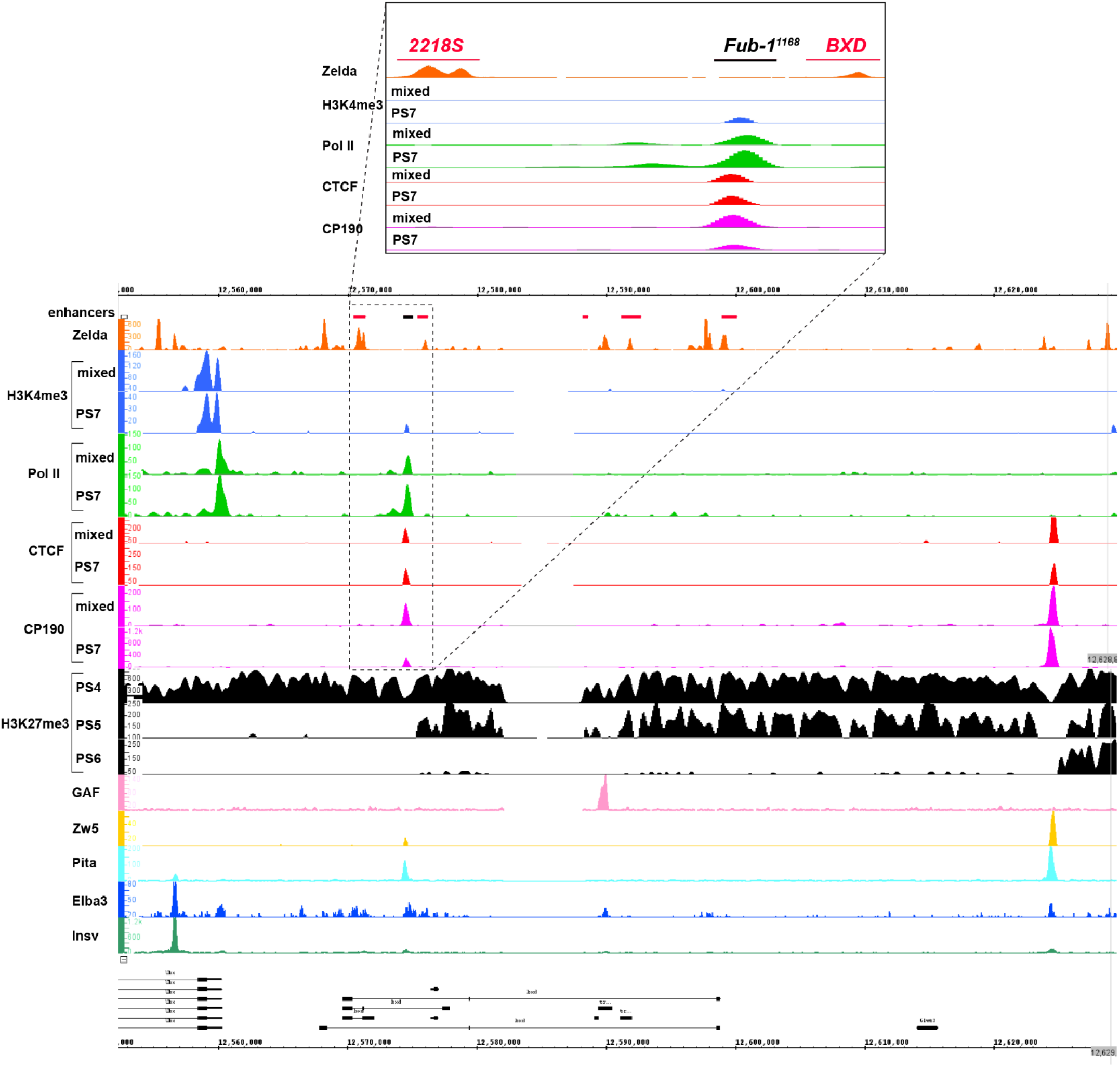
Chip-seq profiles of different chromatin proteins and histone modifications across *bxd/pbx* domain. ChIP-seq profiles for H3K4me3, H3K27me3, Pol II, CTCF, CP190 ^1^, Zw5, Pita ^2^, Elba3, Insv ^3^, Zld ^4^, GAF ^5^. The putative enhancers are indicated by a pink bar below coordinate line.

**Figure S3.**
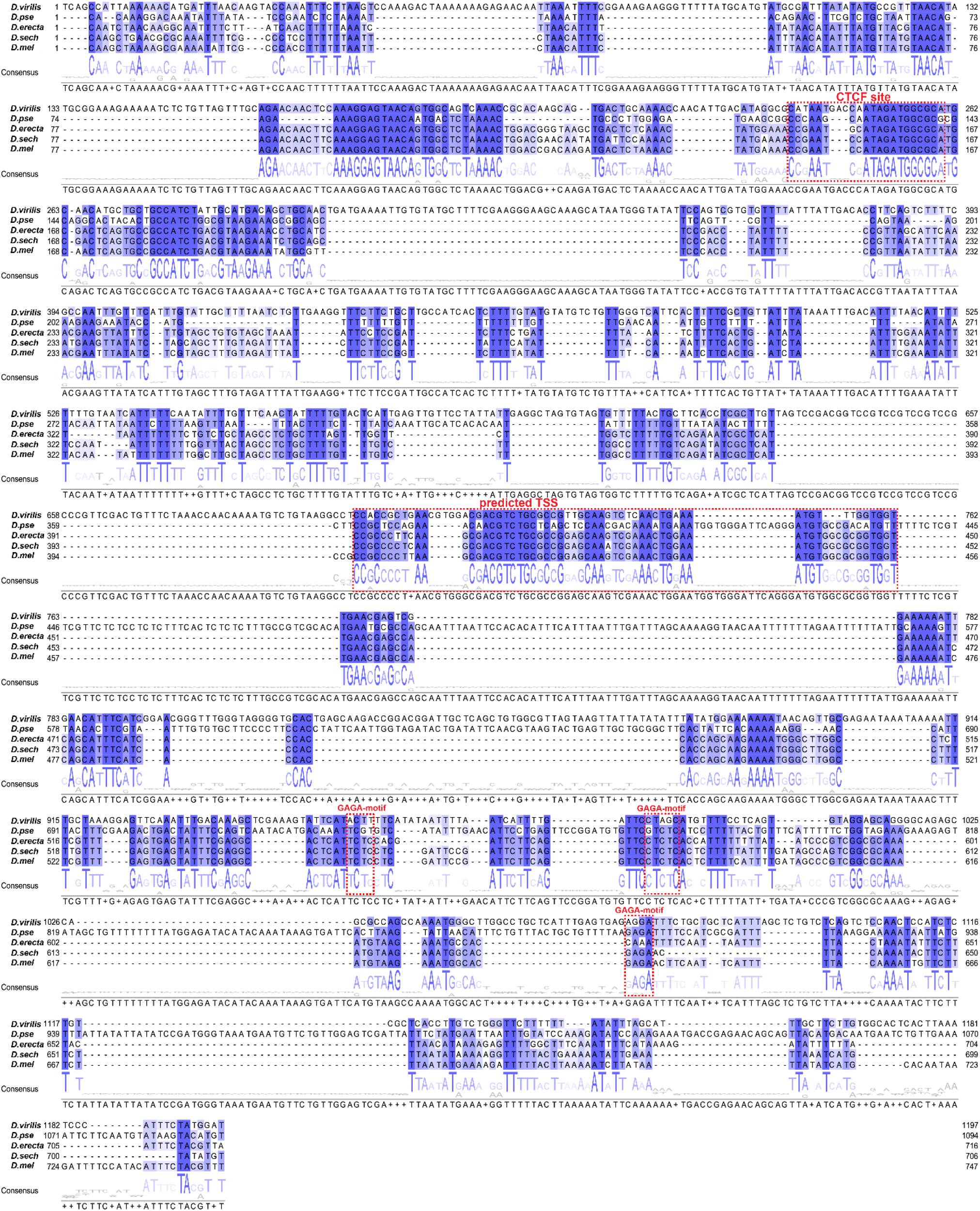
*Fub-1* sequence conservation. Sequence comparison of *Fub-1* boundaries from *D. virilis, D. pseudoobscura, D. erecta, D. sechellia* and *D. melanogaster*. The *Fub-1* sequences of *D. melanogaster* and four other *Drosophila* species were assembled, and stacked up with the ClustalO program to align the high homology region conserved among the five species. Sequences that are homologous in all four *Drosophila* species are highlighted in shades of blue representing degree of conservation. The dCTCF recognition sequence in *HS1*, GAGA-motifs and predicted TSS are enclosed in red rectangles.

**Figure S4.**
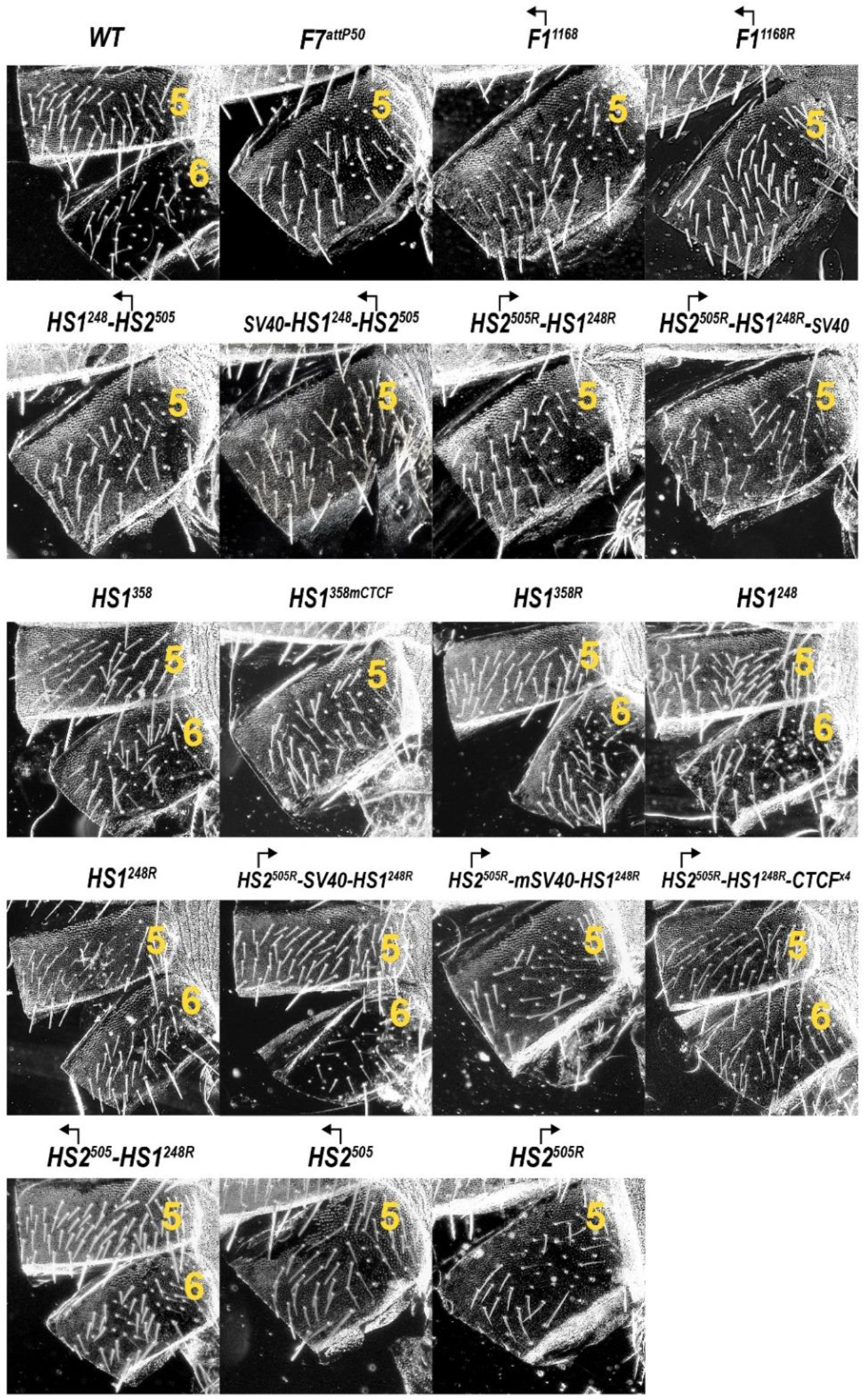
Morphology of the abdominal segments (numbered) in males carrying different variants of *Fub-1* in *Fab-7^attP50^* platform in the dark field.

**Figure S5.**
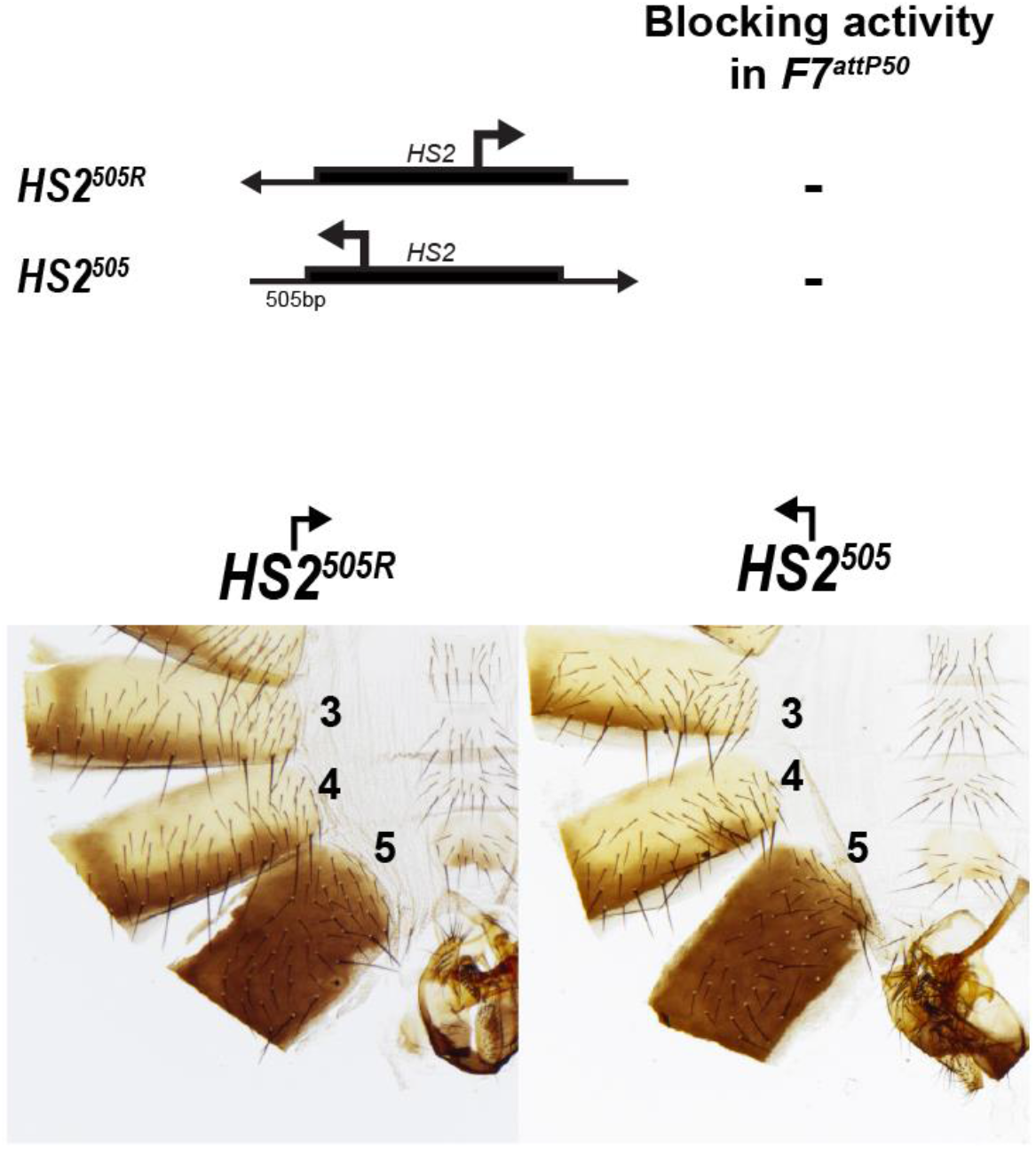
Morphology of the abdominal segments (numbered) in males carrying *HS2^505^* and *HS2^505R^* fragments in *Fab-7^attP50^*.

## Notes

### Competing Interest Statement

The authors have declared no competing interest.

